# Mechanical evolution of DNA double-strand breaks in the nucleosome

**DOI:** 10.1101/254680

**Authors:** Fabrizio Cleri, Fabio Landuzzi, Ralf Blossey

## Abstract

Double strand breaks in the DNA backbone are the most lethal type of defect that can be induced in the cell nucleus by chemical and radiation treatments of cancer. However, little is known about the potentially large differences in the outcomes of damage between free and nucleosomal DNA, leading to corresponding differences in damage repair capability. We performed microsecond-length molecular dynamics computer simulations of nucleosomes including double-strand breaks (DSB) at various sites, to characterize the early stages of the evolution of this important DNA lesion right after its formation. We find that all DSB configurations tend to remain compact, with only the terminal bases interacting with histone proteins; the interacting molecular structures are studied by looking at the essential dynamics of the relevant DNA and histone fragments, and compared to the intact nucleosome, thus exposing key features of the interactions. Moreover, we show that the broken DNA ends at the DSB must overcome a free-energy barrier to detach from the nucleosome core, as measured by means of umbrella sampling of the potential of mean force. Finally, by using state-of-the-art calculation of the covariant mechanical stress at the molecular scale, we demonstrate that, depending on the DNA-core separation distance, the coupled bending and torsional stress stored in the detached DNA can force the free end to either stick back to the nucleosome core surface, or to open up straight, thus making it accessible to damage signalization proteins.

## Introduction

Double-strand breaks (DSB) in the double helix of the DNA molecule are defined as the cleavage of the phosphate-sugar backbone on both sides, the two cuts being comprised within 10 base pairs (bp) at most. Such an occurrence is only one among the many types of DNA lesions that a cell suffers at any time,^1–3^ aside of single-strand breaks (SSB), base loss (AP site, removal of one purine or pyrimidine), base cross-linking or dimerization (such as cyclobutane pyrimidine dimers and 6,4 photoproducts), various types of various oxidative defects by reactive oxygen radicals (e.g., oxidation of guanine to 8-oxoguanine, which substitutes for thymine in the replication, or oxidative deamination of cytosine, which becomes uracil). However, although being much less probable than most other types of lesions, and with relatively fast repair kinetics,^4^ DSBs stand out as the most critical lesions to the DNA, since they ultimately lead to chromosome breakage and genome instability, cell mutation, or apoptosis.^5^ Moreover, several simpler lesions of the type mentioned above, can evolve into SSBs and DSBs upon further chemical processing, both during the repair process, and because of the interaction with other major nuclear proteins.

Because of their cytotoxic effectiveness, inducing DSBs in the DNA of malignant cells is one of the major objectives of chemo‐ and radiotherapy of cancer. Many powerful antitumor antibiotics, such as the enediyne C-1027, abstract hydrogen atoms at several C′ sites in the backbone ribose, initiating an oxidative chain that leads firstly to SSBs (mainly at adenylate and thymidylate residues), and then to DSBs, with two cleaved residues at a distance of 1-2 bp, often sequence-specific.^6^ Neocarzinostatin (NCS), produces 2-deoxyribonolactone (L) as part of a bistranded AP lesion, in which the complementary strand is directly cleaved within 2-3 nucleotides of L.^7^ High-energy radiation creates swarms of ionization products, both directly on the DNA structure and, most importantly, on the surrounding water molecules.^8^ The free radicals produced in the process can attack the backbone and induce many different lesions, often clustered over short distances. Notably, DSBs are produced by ionizing radiation with a relatively high probability, and the terminations at 5′ and 3′ strand ends are typically more complex than for DSBs produced by enzymatic cutting, making their repair more complex and error prone.^4, 9^ Repair of radiation-induced DSBs by both homologous recombination (HR) and non-homologous end joining (NHEJ), requires extra steps prior to the rejoining, to treat and excise the ”wrong” chemical terminations, by implicating very large proteins such as PK (500 kDa and about 16 nm wide, much larger than a single nucleosome) and spreading of damage signals over hundreds or thousands of bases around the lesion, such as phosphorylation of *γ*-H2A.X histone. Moreover, while isolated damage sites, such as AP, SSB, L, are efficiently incised during base excision repair, clustered lesions are much less efficiently repaired. For example,^10^ it was shown that a 8-oxoguanine and an AP-site present within a cluster are processed sequentially, limiting the subsequent formation of DSBs to < 4%; by contrast, when two AP-sites are contained within the clustered DNA damage site, both AP-sites are incised simultaneously, giving rise with high probability to a DSB.

The DNA of eukaryotic genomes is packaged into arrays of nucleosomes, which appear as spheroidal particles that make up the chromatin structure. About three quarters of the total nuclear DNA are included in the nucleosomes, with the remaining DNA acting as ”linker”, in a sort of beads-on-a-string assembly. The nucleosome core particle consists of 147 bp of DNA tightly wrapped around a histone protein octamer containing two copies of four histone proteins (H2A, H2B, H3, and H4). The lysine-rich N-terminal tails of the histones extend from the protein core, making various contacts with the DNA minor groove (see Figure 1 below).

**Figure 1.**
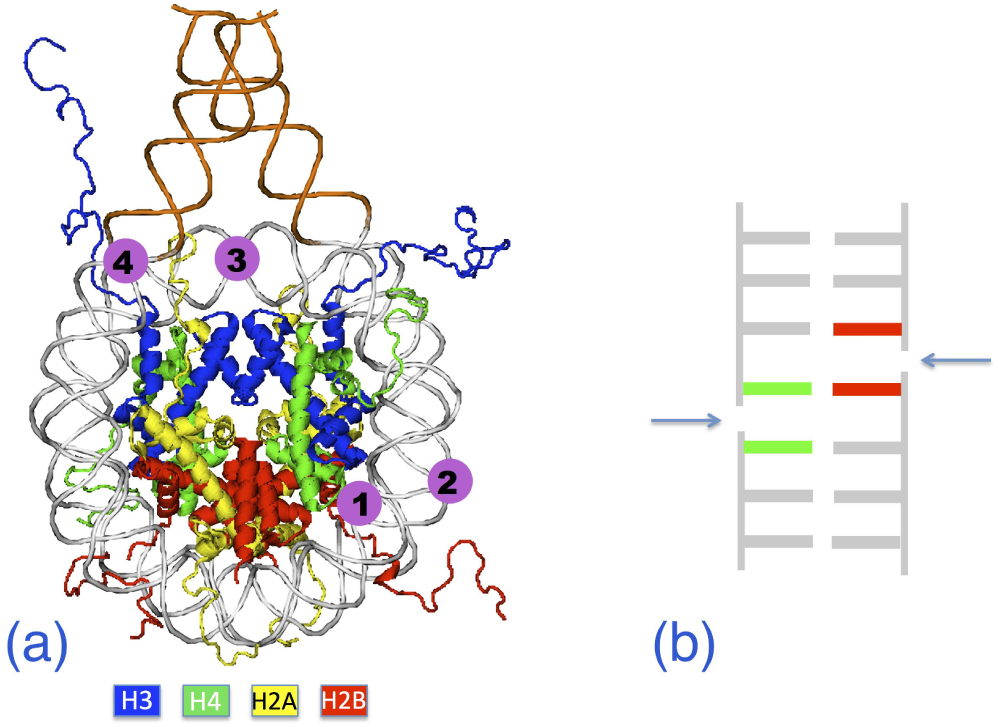
**(a)** Schematic of the nucleosome from experimental structure 1kx5. DNA is shown in grey tube representation; in orange, the 20 bp added to the experimental structure on each end. Histone pairs H3, H4, H2A, H2B are in cartoon representation, color-coded as shown below. Positions numbered 1 to 4 indicate the insertion sites of double-strand breaks (DSB). **(b)** Schematic structure of the DSB spaced by 1 bp, in each of the M1-M4 models. Arrows indicate the site of cut on each backbone; the green-red central base pair is an A…T for all models..

How the complex structural environment of chromatin is altered in the presence of DNA lesions is a longstanding question in the study of the cellular response to DNA damage.^11, 12^ Incorporating a lesion within a nucleosome core particle introduces several features not present in naked (linker) DNA, which affect its reactivity. Firstly, wrapping the DNA around the octameric core introduces mechanical heterogeneity into the duplex, resulting in regions that are bent and/or in which base stacking is deformed. Secondly, the large number of lysine (Lys) and arginine (Arg) residues present in histone proteins (more than 22% on average, Lys being especially abundant in H2B, where it represents 16% of the total) can directly interact with the lesion; in particular, Lys side chains are directly involved in AP cleavage within nucleosomes, via Schiff base formation.^13^ Furthermore, the whole nucleosome structure becomes less compact: histones at DSBs are susceptible to extraction in low salt,^14^ implying a weaker interaction between DNA and histones at DSBs; and biophysical studies demonstrate that DSBs lead to a localized chromatin expansion at DSBs.^11^

Computer simulations are increasingly demonstrating their utility, by allowing deeper analysis of experimental data, also in conditions where experiments are difficult to carry out. Molecular dynamics (MD) simulations of nucleic acids are today capable of following the system evolution over length and time scales approaching the real experimental set up.^15, 16^ The main difficulties lie in the appropriate treatment of reactive configurations with such molecular databases as AMBER or CHARMM, which were originally conceived and adjusted on the physical and chemical properties of small fragments of perfect DNA or RNA. For this reason, the domain of molecular lesions to nucleic acids and chromatin by molecular simulation methods remains still little explored. We recently completed a first molecular dynamics study of the mechanical evolution of SSBs and DSBs in random-sequence DNA oligomers.^17^ A DNA fragment of length 31 bp with B conformation, held at the two extremities by fluctuating springs, was taken as representative of the exposed ”linker” DNA between two nucleosomes in the chromatin fiber. We studied the mechanical response under tensile force of SSBs and DSBs with different spacing between the two strand cuts (or “nicks”), by means of MD simulations mimicking single-molecule force spectroscopy experiments. The results indicated that the absolute values of force necessary to break up a DSB-damaged, free DNA fragment can be very large, of the order of 100 pN, at elongations of ∼20%. Such values of longitudinal stress and strain are unlikely to be observed in the normal dynamics of chromatin, nor during chromosome mitosis. Most importantly, however, that study demonstrated that thermal fluctuations are unable to provide the energy necessary to overcome the barrier to rupture, unless the two DSB cuts are separated by 2-3 base pairs at maximum.

In the present work we turn to investigating the mechanical evolution of DSBs in a nucleosome immediately after the backbone breaking event, by using very-large-scale molecular dynamics simulations in the microsecond time scale. All-atom MD simulations of the nucleosome have started a few years ago, initially restricted to a 10-100 ns time scale,^18, 19^ and very recently extended to the microsecond time scale;^20, 21^ these works provided already a substantial description of many special features of DNA wrapped in a nucleosome, such as the effects of added torsion and bending, histone-DNA contacts, and much other. However, the structure and dynamics of DNA defects of any kind are yet unexplored, in the much wider context of the nucleosome. Here, we search for specific mechanical signatures induced by the DSB, by simulating microsecond-long trajectories of the entire system, embedded in a large box of water and neutralized with point ions. We use different analysis methods to characterize the mechanistic aspects of DSB structural evolution, at various positions in the nucleosome. Firstly, we look at the long-wavelength thermal fluctuations of the system, extracting the essential dynamics from the covariance matrix of the atomic displacements around the DSB region. Secondly, we determine the lability of the broken-DNA adhesion to the histone octamer, by force-pulling with the ”umbrella sampling” method. Finally, we characterize the role of mechanical bending and torsional stress, in determining the evolution of the broken DNA ends at longer times.

Altogether, these analyses allow to trace a mechanical path of the evolution of individual strand breaks into a fully-developed DSB, up to the final fracture event, as well as suggesting the most probable late-stage mechanical evolution of the damaged nucleosome. We conclude that DSBs can be resistant to spontaneous disassembly by thermal forces, thanks to the strong DNA-protein interaction within the nucleosome; in fact, mechanical forces of some importance are needed to open up the DNA structure at the break site, in a manner that appears to require the intervention of external agents; once the DNA structure starts to be opened, its fate depends on the amplitude of the displacement, the DNA being able to fold back to its original configuration, or to straighten out from the nucleosome core for a large enough initial opening; also in this case, substantial external forces are necessary to bring the DNA ends to a sufficiently wide opening. These crucial findings should have profound implications for the early stages of DNA damage detection and repair, for example implying that damage marker proteins (such as Ku70/80, which interacts strongly with broken DNA ends), should also be capable of exerting complex mechanical actions, for the damaged DNA to be accessible to the repair agents subsequently recruited in cascade.

## Materials and Methods

### Molecular structures of damaged nucleosome

We obtained the nucleosome molecular configuration from the RSCB Protein Database, entry 1kx5.^22^ This is an x-ray structure of the entire histone octamer with 147 DNA bp resolved at an average RMS of 1.94 Å, reconstituted from human nuclear extract expressed in *E. coli;* only 6 histone residues were unidentified in this experimental structure, with respect to the known histone sequences, therefore the model can be considered nearly complete. The 147 bp DNA is a palindromic sequence, chosen to maximize the degree of ordering and increase the x-ray spatial resolution. To obtain a model structure useful for our computer simulations, we removed all the crystallization water molecules and ions from the published structure, and added two DNA extensions of length 20 bp at each end of the nucleosomal DNA, with repeated sequence d(AGTC).^20^ DNA bases are numbered from 1 to 187 in each chain, one running clockwise and the other counter-clockwise, the dyad being located at basis 94 of each chain. This pristine nucleosome model without strand breaks is shown in Figure 1a, and will be labelled O in the foregoing.

DNA is wrapped left-handed about the histone core, making two nearly complete turns that join at the dyad symmetry point; the two DNA turns define two circles lying in two ideally parallel planes, with a superhelical symmetry axis perpendicular to the center of the circles (for a thorough discussion of nucleosome geometry and structure, see e.g. Ref.^23^). The relaxed DNA double helix makes a complete twist around its double-helical axis, about every 10.4 bp, defining a major and a minor groove; therefore, when turning around the histone core, the wrapped DNA makes 14 nearly full twists. Correspondingly, 14 contact points between DNA and proteins can be identified within the nucleosome structure, loosely situated at the minor groove locations facing inwards.

Based on these geometrical features, we defined 4 potentially interesting sites along this wrapped structure, where to place a DSB in a ”mechanically significant” position, labelled 1 to 4 in Fig 1a. Correspondingly, we introduced a DSB at an inner contact site (model M1); at an outer non-contact site (M2); at the dyad (M3); and at the entry point of the nucleosome (M4). To create a DSB at each such locations, we introduced 5′-OH and 3′-phosphate terminations at each end of the break, respectively between^1^: bases C69-T68… A120-G121 in M1; bases C73-A74… T114-T113 in M2; bases A94-T95… A94-T95 of both chains in M3; bases T22-A21… T167-G168 in M4. In this way, the two backbone cuts of each DSB are spaced by 1 bp always comprising an A…T pair (Fig 1b), which remains initially bonded by only its two hydrogen bonds, plus the stacking interactions on each intact side of the chain, while the other half of stacking is readily reduced, as soon as the MD relaxation starts.

The CHARMM-27 force field database^24, 25^ and its extension to treat nucleic acids^26^, ^27^ were used for the molecular bonding and non-bonding force parameters. Strict comparisons between CHARMM-27 and AMBER force fields^28, 29^ ensure that the results of long-time, finite-temperature MD trajectories of nucleic acid fragments with largely different conformations are consistent, and able to correctly reproduce the key structural quantities (bond angles, hydrogen-bond structure, base tilt, twist, shuffle, etc.) compared to experimental data. However, in all cases great care must be taken by performing sufficiently long preparatory annealing cycles of the water and ion background, while keeping the nucleosome still, to obtain the right water density and allow a realistic arrangement of the counter-ions around the phosphate backbone, prior to starting the microsecond production runs.

### Molecular dynamics simulations

For the molecular dynamics (MD) simulations we used the GRO-MACS 5.1 computer code.^30, 31^ Nucleosome models O and M1-M4 were solvated in water box of size 14.5 or 18×19×10 nm^3^ with periodic boundary conditions in the three directions, containing about 82,600 or 110,500 TIP3P water molecules, plus 480 Na+ and 250 Cl^−^ ions to ensure neutralization of the phosphate backbone charge, and a physiological salt concentration around 0.15 M. All the MD simulations were carried out at the temperature of 310 K and pressure of 1 atm, or 350 K and 50 atm for the thermal stability study. Because of the requirements of stress calculations (see below), we could not use standard Ewald-sum electrostatics but plain cut-off Coulomb forces. This is known to be at the origin of possible artifacts, therefore we adopted for both electrostatics and long-range non-bonding forces an unusually large cut off radius of 1.6 nm. The DNA terminal ends (linker) were restrained by soft harmonic constraints, allowing a fluctuation of ±5 Å, to represent embedding in the chromatin structure. We used rigid bonds for the water molecules, which allowed to push the time step to 2 fs for the thermal equilibration runs, and to 1 fs for the force-pulling simulations. Typical preparatory constant-*{NPT*} MD runs lasted between 10 and 20 ns; force-pulling simulations were carried out for 10 ns, and the subsequent force-free relaxation lasted up to 400 ns; thermal stability simulations at constant-{*NVT*} extended to ∼1,000 ns for O and M2-M4, and up to 1,800 ns for the M1 model. Overall, the study used about 4.2 million hours of CPU time on 2048 IBM BlueGeneQ processors (IDRIS supercomputing center in Orsay), and about 800,000 hours on 896/1064 Broadwell Intel E5-2690 multi-core processors (CINES supercomputing center in Montpellier), with typical running times of 1.3 and 7 ns/hour on the IBM and Intel machine, respectively. About 1.5 Terabytes of raw data were accumulated over a period of 8 months, from March to October 2017, for subsequent post-processing. All-atom microsecond trajectories for the nucleosome (DNA+proteins, minus water and ions), stored in the compact GROMACS-xtc format at frame intervals of 40 ps, are freely available upon request to the authors.

### Steered MD and umbrella sampling

Steered molecular dynamics (SMD) was performed on the fragments with the constant-force pull code available in GROMACS, only on the M1 model. In this case, we enlarged the water box to 18 nm in the x-direction, to allow possible outward extension of the broken DNA end, resulting in a system of 107,000 water molecules. Since the objective was to promote the detachment of one of the broken DSB ends from the nucleosome core, we applied a constant force parallel to the direction x and perpendicular to the superhelical axis, by means of a harmonic-spring fictitious potential attached to the C4′ and P atoms of the last two base pairs at one DSB end. After some tests, the spring constants were set at 100 and 75 kJ mol^−1^ nm^−2^, respectively for the two DNA strand ends farther and closer to the nucleosome surface. To provide a reaction force keeping the system in place, all the atoms of the H3 opposite to the DSB were retained by soft harmonic restraints, with a spring constant of 250 kJ mol^−1^ nm^−2^. Pulling speeds in the range 1 to 5 mm/s were used, with most SMD simulations being carried out at the lowest speed. Forces and displacements were recorded at intervals of 5-10 time steps. Umbrella sampling was performed by extracting 100 configurations spaced by 50 ps during the first 5 ns of the force pulling simulation; force bias was progressively reduced from 100 down to 10 kJ mol^−1^ nm^−2^, to extract the zero-bias limit of the free-energy profile; the weighted-histogram analysis was used to interpolate and connect the data from discrete configurations.

### Molecular stress calculation

We use the so-called covariant central-force decomposition scheme (CCFD,^32, 33^) for the intra‐ and intermolecular forces, which ensures conservation of linear and angular momentum of the molecular systems under very general conditions. The method is implemented in a special-purpose patch to GROMACS 4.6, which reads (all or part of) a MD trajectory for the selected subset of atoms for which stress is to be computed, and performs the entire analysis. Since the patch (called GROMACS-LS by the authors^32, 33^) constrains the code to run in serial rather than in parallel, care must be taken to define properly the subset of interest in order to avoid prohibitive computing times. We prepared simple scripts to extract the principal components of the stress, compare stress fields from different simulations, and write the outputs in the portable Gaussian-cube format for visualization. Comparison between stress fields from different MD runs poses an extra care, since the structures need to share exactly the same box size and center, to avoid numerical artefacts from the cancellation between large positive and negative values. According to the CCFD scheme, stress fields are calculated by GROMACS-LS on a continuous grid superposed on the molecular structure; however, stress components and individual force contributions (pair, angle, dihedral, etc.) can also be projected back on the atom sites by defining a conventional (but non unique) atomic volume.

## Results

### DSB dynamics at different nucleosome positions

In our previous study on linker DNA fragments,^17^ it was found that DSB can be very stable against thermal fluctuations, unless the two cuts on the backbone are very closely spaced. In particular, we obtained an average bond lifetime of the order of 50 ns at T=350 K for the DSB with a single-bp A… T pair, and from these data we extrapolated lifetimes of the order of hundreds of milliseconds for a DSB with 2-bp spacing, and up to several hours for a DSB with 3-bp spacing. Part of that study was conducted at relatively high temperatures, in order to accelerate the rare event of bond rupture, however allowing a meaningful extrapolation down to physiological temperature.

Based on such results, we decided to use the most favorable DSB configuration in the present study, in order to increase the probability of eventually observing DNA break up. Therefore, we introduced in all models M1-M4 one single-bp DSB with a central A… T, which is the weakest bonded bp. We ran the MD simulations at the temperature of T=350 K, or about 77° C, in order to stimulate the thermal dynamics of the system, while remaining within a range of vibrational excitations that is still meaningful for the molecular force field used. MD trajectories were extended to ∼1 μs for the M2-M4 models, and up to 1.8 μs for the M1, which displayed some potentially more interesting dynamic features. The reference model O with the intact nucleosome was simulated over a shorter trajectory of 500 ns. Shorter MD trajectories were also run at T=310 K for all models, for comparison.

We firstly present the results for the models M2-M4. For all these three models, we could not observe any substantial evolution of the DSB into a fully broken DNA, over the whole duration of the simulation, despite the relatively high temperature. While it cannot be excluded that such an event could be produced over longer times, this is an increase of more than a factor of 20 in lifetime compared to the free (linker) DNA; once scaled down to 310 K, this translates in spontaneous DSB dissociation times at least in the 100-μs time scale, or longer, for the most favorable (i.e., least bound) DSB configuration, which represents therefore a lower bound for the dissociation time. Some representative snapshots of the thermalized molecular structures at the DSB site are displayed in Suppl. Fig. 1.

The bonding configuration of the central base pair remains on average rather close to that of the pristine nucleosome, with the H-bonds providing a large fraction of the cohesive energy, and the mildly deformed stacking ensuring a substantial structure stability. An example can be observed in Figure 2, in which the time evolution of the H-bond lengths for the central A… T bp of model M3 are shown. The three bonds formed by the N1 (adenine), O1 and O2 (thymine) donors are indicated in red, blue and black, respectively. The relative strength of individual H-bonds in the A… T bp can be theoretically estimated^34^ to be about 10:4:1 for the N1:O1:O2. The last one is not usually accounted as a true H-bond, since it is very weak and with a length fluctuating around 2.8 Å. Indeed, in our MD simulation the central N1 bond remains always in the range [1.8-2.1] Å RMS (note that the simulation is at high temperature); the side O1 is more dynamic than the corresponding bond in normal DNA, with an average length of 2.3 Å (∼2.05 in normal DNA), and quite large RMS fluctuations due to the larger rotational freedom of the DSB about the central axis; the O2 length remains well beyond the definition of H-bond, fluctuating about an average of 3.2 Å. Overall, these interactions provide enough bonding to keep the DSB in place, even in this M3-dyad position that is the farthest from the histone protein core, among all the DSB configurations studied.

**Figure 2.**
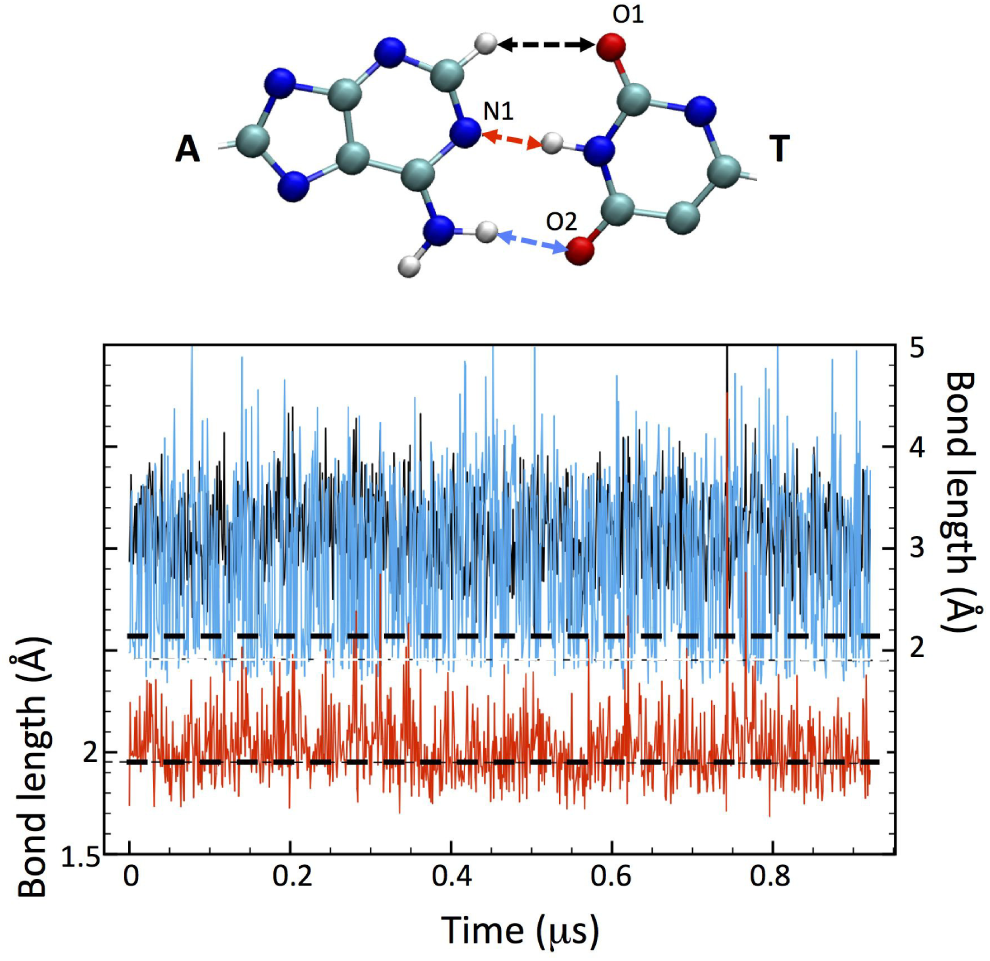
Plot of the H-bond length for the central A… T bp of the model M3, for a MD simulation of 950 ns at T=350 K. Red, blue and black traces relative to the respective arrows in the scheme above, with the A…T bp in ball-stick representation. The red arrow indicates the central H-bond with adenine N1, the blue arrow the side H-bond with thymine O1, and the black arrow the (very weak) H-bond with thymine O2. Horizontal dashed lines in the plot indicate the reference, T=350K H-bond distances of 1.9 (lower) and 2.1 Å (upper).

To characterize the dynamic motion of the DNA and of the closest protein residues around the DSB region, we performed for each model a study of the essential dynamics.^35, 36^ This method of analysis looks at a small subset of collective coordinates of the system, to extract the large-scale, anharmonic movements (bending, torsion, etc.) that dominate the global molecular dynamics. Such large-scale motions are deduced from the eigenvectors of the displacement covariance matrix:

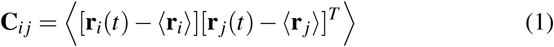

where **r**(*t*) is the vector of time trajectory of each atom *i,* and 〈…〉 indicates time-averaging. The symmetric matrix **C** is of rank *3N* for a set on N atoms, and can be diagonalised by standard methods as **C** = **V**Λ**V**^*T*^, with Λ the diagonal matrix of the eigenvalues and **V** the matrix whose columns are the eigenvectors. Quite often, only the first few eigenvalues represent the largest fraction of the total weight, so that by looking at the first few eigenvectors, the large-scale, long-wavelength motion of the system can be appreciated. Notably, this type of analysis makes no assumption about harmonicity of displacements from equilibrium, therefore it can identify large conformational fluctuations. The remaining eigenvectors (that is, nearly all) describe the harmonic motions, and could be linked to the vibrational spectrum of the molecular system.

We firstly perform the analysis for the regions surrounding each location M1-M4 in the pristine nucleosome, model O. Typically, the analysis is restricted to a length of about 7 DNA bp on each side of the DSB, plus the 15-20 histone residues in the closest neighborhood of the DSB. MD trajectories are sampled at a rate of 40 ps^-1^. This first analysis of the undamaged system provides a spectrum of eigenvalues, from which we extract the first few significant ones, and an average reference configuration for each M1-M4 site. Then, we repeat the same analysis on each of the independent trajectories including a DSB at the M1-M4 positions, by using as reference molecular structure the corresponding average from model O, so as to highlight deviations from the normal DNA dynamics.

A key quantity providing information about the large-scale (or ”long-wavelength”) movements of the fragments implicated in the DSB comes from the study of the first few eigenvectors, and of their root-mean-squared fluctuation (RMSF) on a atom-by-atom basis. These new atomic variables capture the contribution of each group of atoms to the principal collective movements, as filtered out by the most important eigenvectors. For all the M1-M4 models, the first 4 eigenvectors are found to cover 65% of the weight, the 5-15 ones are responsible for another 20%, and all the remaining 3*N*-15 for the last ∼15%. Such a distribution is less extreme for the O model, in which large-scale movements are quite more restricted, with the first 15 eigenvalues carrying about 55% of the total weight. The physical meaning of such principal eigenvectors can be appreciated, for example, with the timeframe plot of Suppl. Fig. 3, where the extreme configurations spanned by the large-scale motion of the first eigenvector, for the DNA fragments in models M1 and M3, are all simultaneously represented; the frames are colored from blue to red, the ordering showing how each atom’s motion spans between the extremes of the eigenvector. It can be seen that the principal eigenvector for M3 describes quite homogeneous, local fluctuations of all DNA bases, with just a more evident oscillation along the stacking direction concentrated about the DSB; on the contrary, for the M1 this principal eigenvector describes a dramatic large-scale displacement of the central atoms making up the DSB, which tend to span ample areas across orthogonal planes, by turning about the backbone. This largely different behavior between M1 and the other models M2-M4 is discussed further in the following.

In Figure 3 we plot the RMSF for the first 4 eigenvectors of each DSB model; each plot compares the RMSF for the fragment of DNA surrounding the DSB (black lines), with the corresponding RMSF of the same fragment intact (red lines). For the M2-M4 models, it can be clearly seen that the RMSF of the DSB fragments is comparable to that of the same fragment in the reference model O; despite local quantitative variations, also of some importance between the various DNA bases, the black and red traces remain always close to each other, for each eigenvector, within a range of 0.1 in the arbitrary units of the RMSF. Moreover, the regions of the DSB and the base-pairs immediately adjacent (indicated by grey shaded areas) do not seem to display a peculiar or specific behavior, compared to the bp more distant from the DSB locations. Only the 1st and 3rd eigenvectors of M4 are somewhat outstanding compared to all the others, since they display a curiously even, or less random, distribution of displacements among all the bp. As it can be seen in the detailed eigenvector plots in Suppl. Fig 4, this coordinated motion correspond to an ample twisting about the main axis, which exists both for the O and M4 model, therefore independently on the presence of the DSB. Notably, the M4 location is the most ”straight” and less perturbed, compared to the rest of the strongly curved and bent DNA in the nucleosome core.

**Figure 3.**
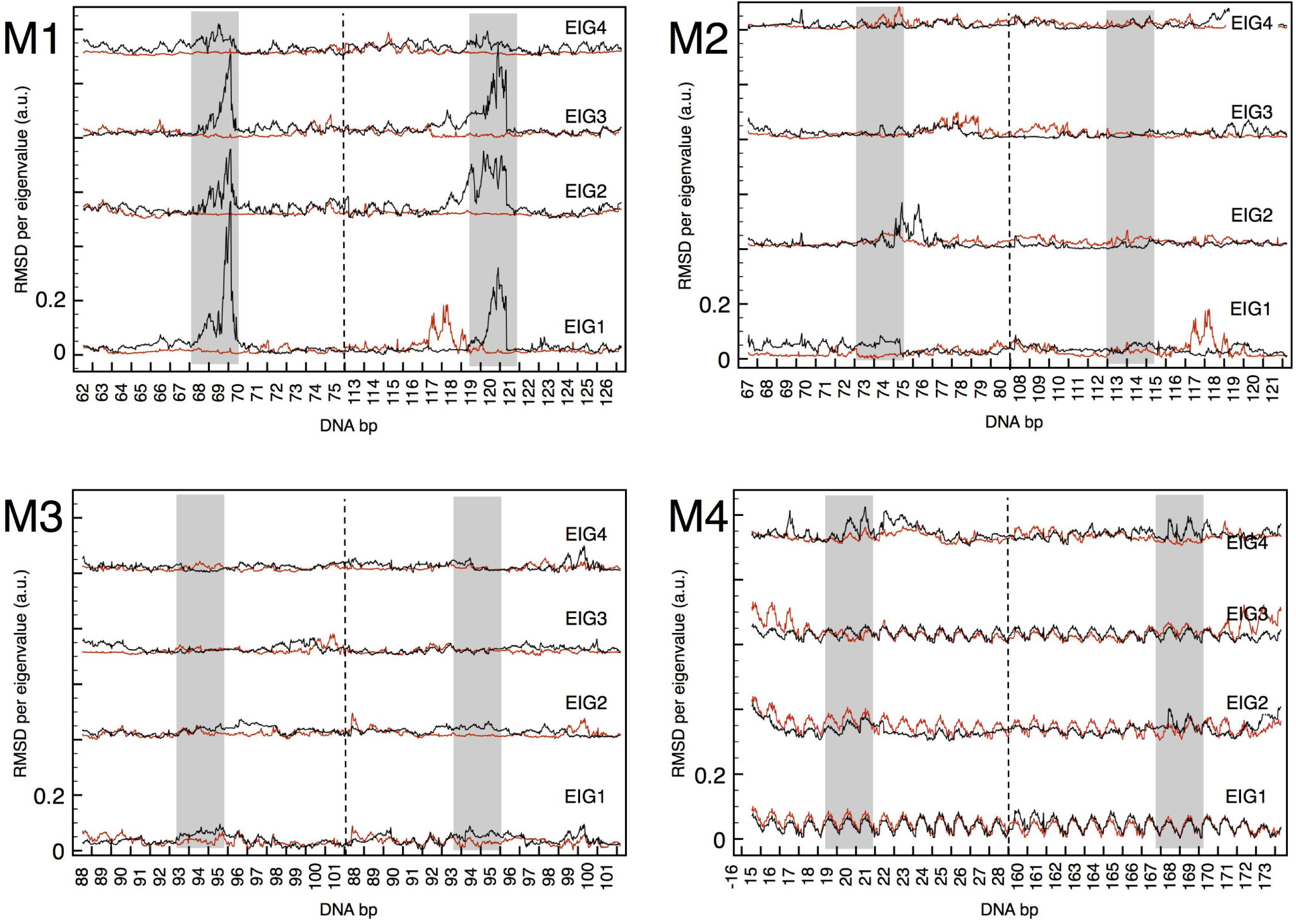
Plot of the RMS fluctuation of the 4 principal eigenvectors, for the DNA fragments including the DSB, of the four models M1-M4. For each eigenvector (respective origins shifted along the ordinate axis) the black line gives the atom-by-atom contribution of the fragment surrounding the DSB, while the red line gives the same quantity for the same fragment intact (from model O). On each abscissa, atoms are grouped by the base number, in two contiguous blocks divided by the dashed line, representing the two parallel strands; the grey shaded regions indicate the position of the DSB (the red-green bases of Fig 1b) for each model.

The same analysis can be carried out for the RMSF of the histone residues closer to the DSB in each model, as shown in Figure 4. Also in this case, for the M2-M4 models it is hard to see a qualitative difference between the data for the intact fragments (red lines), and for the fragments with the DSB inserted (black lines). Both the lysine and arginine residues are overall more mobile than the others, as far as the 4 principal eigenvectors are concerned, describing a dynamic interaction with the DNA. However, with minor variations, this behavior is the same also in the absence of the DSB, therefore it reflects the usual affinity of such residues for the DNA bases. The M3 model is perfectly symmetric, with the two portions of each H3 histone playing exactly the same dynamics, on the two flanks of the DSB. The M1 model, instead, is definitely different, as it was the case also for the DNA analysis in Fig 3 above, and it will be treated later in this Section.

**Figure 4.**
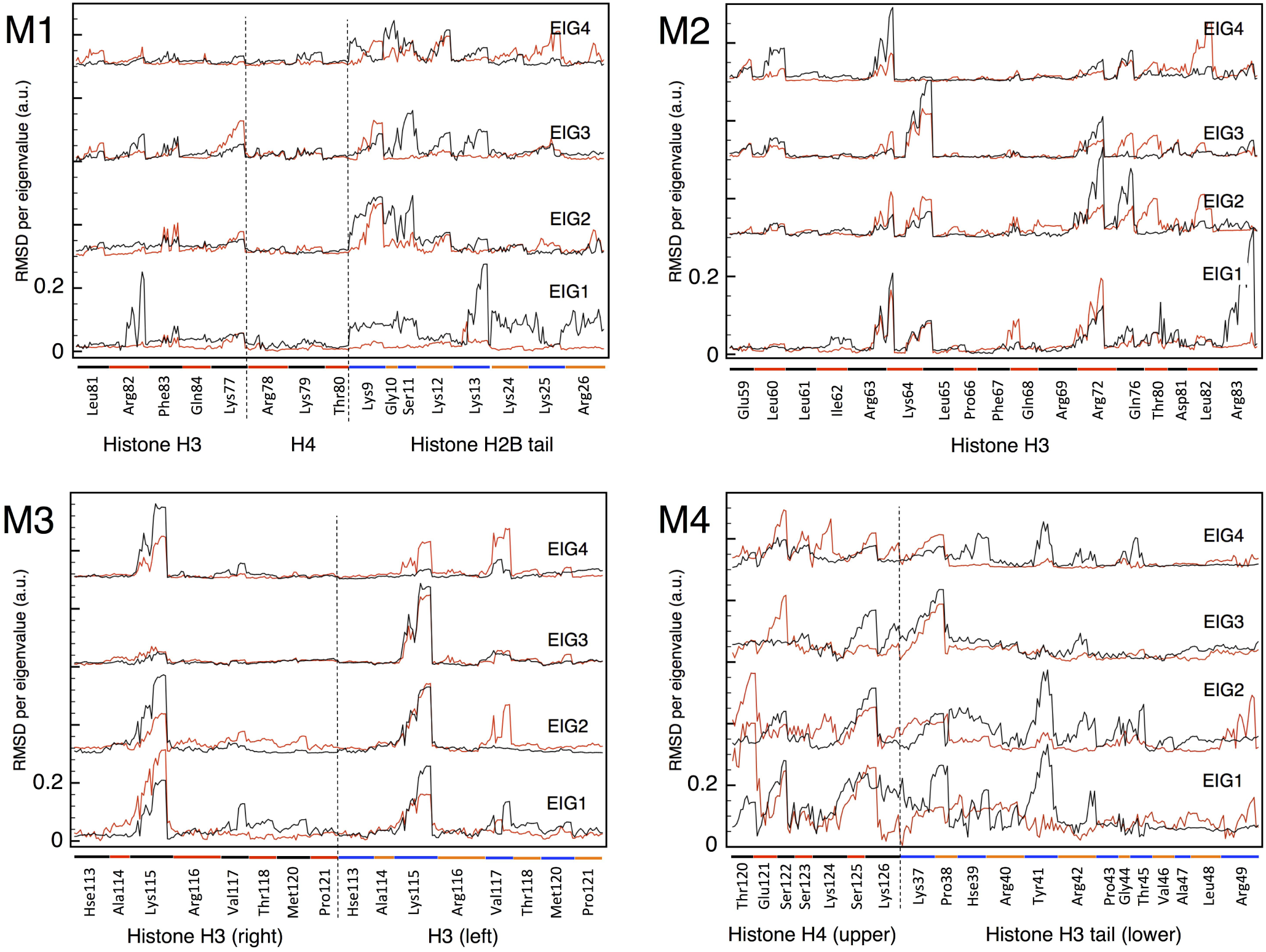
Plot of the RMS fluctuation of the 4 principal eigenvectors, for the histone residues closest to the DSB in each of the four models M1-M4. For each eigenvector (respective origins shifted along the ordinate axis) the black line gives the atom-by-atom contribution of the fragment surrounding the DSB, while the red line gives the same quantity for the same fragment intact (from model O). On each abscissa, atoms are grouped by residues, with a spacing (also indicated by colored bars) proportional to the size (i.e., number of atoms) of each residue.

From the covariance matrix, the Schlitter entropy formula can be used to estimate an upper limit to the excess entropy contribution to the free energy due the presence of the DSB,^37^ as:

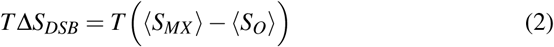

with *MX* = *M*1, …*M*4, and 〈…〉 indicating the time average of the Schlitter entropy for each molecular fragment:

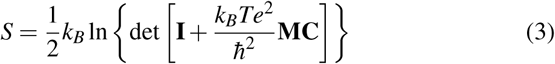

with **I** the identity matrix and **M** the mass matrix, having respectively 1 and the atom masses on their diagonals, and 0 elsewhere. Table 1 reports the values for each DSB model, divided into DNA and histone contribution.

**Table 1.** Upper limit of the excess entropy contribution *T*Δ*S* to the free energy at T=310 K, estimated from the Schlitter formula, Eq 3, for each molecular fragment in the different DSB models.

| DSB configuration | DNA (kcal/mol) | histones (kcal/mol) |
| --- | --- | --- |
| M1 | 89.7 | 29.8 |
| M2 | 25.6 | 6.9 |
| M3 | 38.1 | 2.7 |
| M4 | 47.9 | 5.3 |

The absolute DNA entropy SO from Eq 3 fluctuates about 18±0.5 kcal/mol/K for each base, very homogeneously all along the most part of nucleosome, but increasing to 20 kcal/mol/K in the few terminal base-pairs attaching to the straight segments. If the values of excess entropy of DNA reported in the Table are distributed to the 4 bases (green and red in Fig 1b) comprising the DSB, these correspond to an excess of 35 to 60% for the M2-M4 models, the excess per base being larger in the M4, in agreement with the somewhat larger mobility demonstrated in Fig 3. On the other hand, the excess entropy for the histone residues selected for this analysis remains relatively small, for the three models M2-M4. Despite some difference in the total masses of the groups selected, even when expressed per unit mass instead of per-moles, the absolute entropy of the histones remains comparable, between the model O and the models including the DSB. This is a further conirmation of the relatively minor role played by histone dynamics in the M2-M4 models.

We now turn to describing the behavior of the DSB in the M1 model. Contrary to our expectations, this location in which the DSB is constrained between the histone core and the mobile H2B tail, and close to a DNA-protein contact, was the one to display the most interesting dynamics. The most evident change in the immediate environment of the DSB is the modification of the H2B tail, which can fold into very different interacting positions, starting from the outward extended conformation of the experimental crystallographic structure. This behavior is shown in Figure 5, where the arrangement of the H2B tail is represented for three configurations, averaged over the respective MD trajectories: the reference O model (yellow), the M1 model at T=310 K (cyan), and the M1 model at T=350 K (blue). The low-temperature average configuration of the H2B tail resembles well that of the O model, with the terminal wrapping the minor groove of the DNA strand on the left of the DSB (in the figure); the high-temperature average configuration, instead, has the H2B tail flipped down by about 180 degrees, with the fold of Lys24-25 and Arg26 keeping close contact with the DSB (see the black arrow). That such a configuration may be dynamically sampled over ∼ 1 μs time by only a 40 K temperature difference, means that the corresponding energy barrier (chemical plus deformation) must be relatively small.

**Figure 5.**
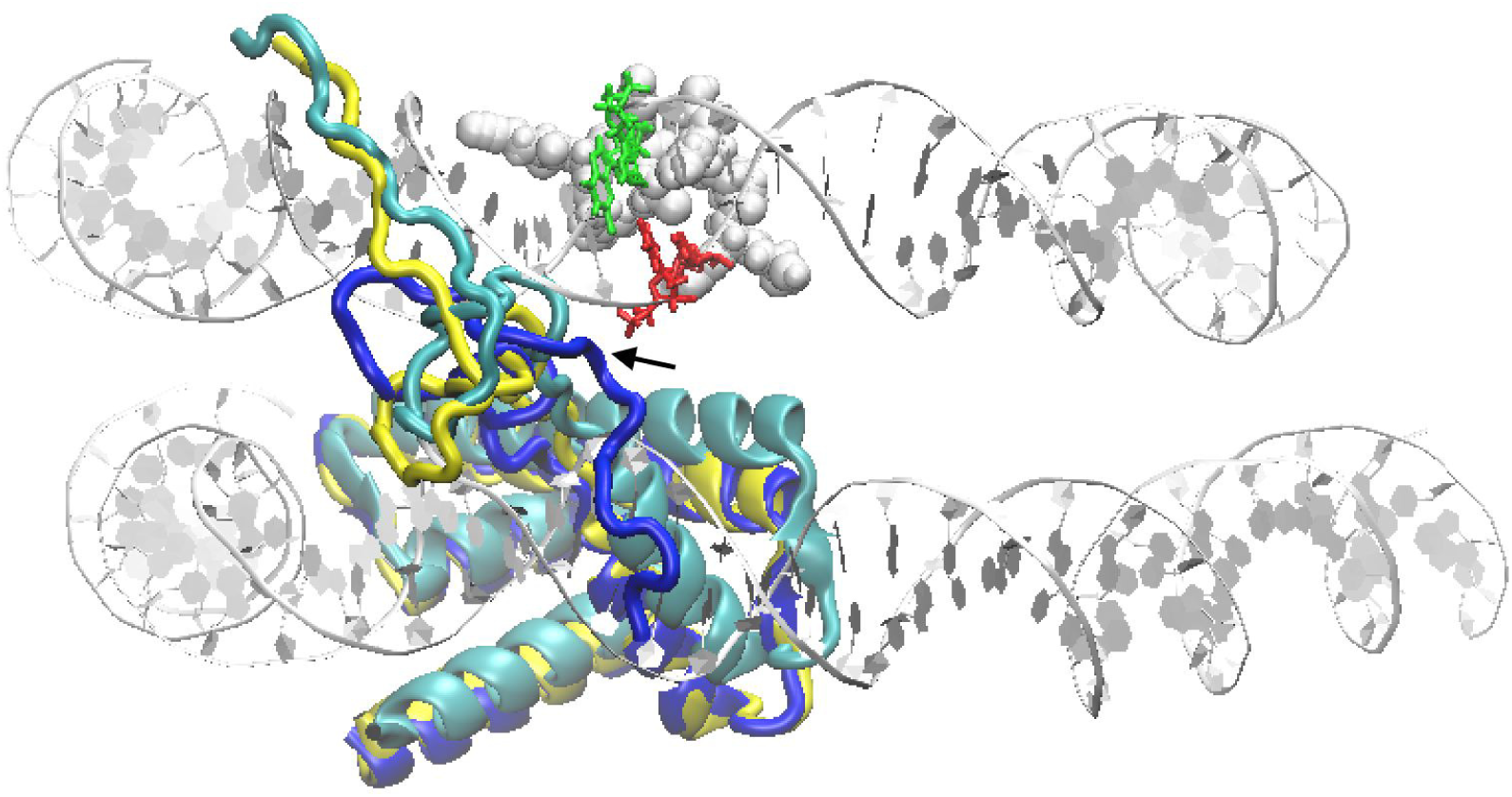
Representation of the time-averaged H2B histone configurations: in the reference O model (yellow ribbons), in the M1 model at T=310 K (cyan) and in the M1 model at T=350 K (blue); the tail of the histone can be seen taking different orientations in the former two, w/r to the last simulation. Two portions of the upper and lower turn of DNA about the histone core are represented in silver ribbons; the spheres on the back represent the H3/H4 residues closer to the DSB; the DSB bases are colored in red-green according to Fig 1b.

In this M1 model, the DSB is constantly enclosed between the two *β*-sheets of H3 and H4, which fluctuate about their equilibrium structure and interact with one side of the DSB, while the H2B tail experiences strong oscillations, coupling with the cut bases of the opposite DSB side. The time evolution of the four bases comprising the DSB (green-red colored in Fig 5) gives a qualitative appraisal of this strong interaction (Suppl. Fig. 2). Notably, the interacting portions of both the two *β*-sheets, and the H2B tail, include more than 60% of lysine and arginine residues, as expected given the strong electrostatic affinity of such amino acids for DNA (notably for G and T,^38^). The DNA ends at the DSB are clearly perturbed by such interactions, and it can no longer be said that the two broken backbones preserve a geometrical continuity, as it was instead observed for the M2-M4 models for the entire duration of the respective MD trajectories.

By looking at the RMS fluctuation of the eigenvalues for the M1 model in Figs.3,4, it can be seen that in this case the group of DNA bases adjacent to the strand breaks take up the majority of the weight, indicative of their participation in the large-scale, ample fluctuations of the open ends of the DSB; also for the histones, it is readily apparent a more dramatic dynamics, especially by the lysine and arginine residues; finally, the values of excess entropies from Table I give a further confirmation of the peculiar large-scale dynamics of this DSB configuration. However, because of these indications, we continued the MD trajectory up to 1.8 μs, but never observed a true mechanical destabilization of the DNA structure: the two broken ends remain firmly in place, even if the two terminal bases on each strand of the broken ends fluctuate quite wildly (see again Suppl. Fig 3, and also the motion indicated by blue arrows in Suppl. Fig 2), while promoting a strong interaction with the protein surfaces.

### Free energy to detach broken DNA ends

As shown in the preceding Section, spontaneous dissociation of one or both DSB ends of a broken DNA from the nucleosome remains a difficult event, never observed in our simulations. DSB opening and detachment of DNA from the nucleosome are likely governed by a free energy barrier of adhesion, which even such critical defect as a fully-cut DNA could not easily overcome, simply by thermal fluctuations. The way to estimate the free-energy barrier in such a large and complex molecular system is to resort to controlled-force pulling, in order to impose the detachment, and then to use the intermediate structures along the reaction coordinate as starting points, for the ”umbrella” sampling of the potential of mean force. From the latter, the free energy barrier along the chosen reaction coordinate can be extracted.

As briefly described above, we used as reaction coordinate *ζ* the separation distance between the moving DSB end and the histone core surface. This was measured by taking the center of the DNA axis, at the average position of the C4′ and P atoms of the last two base pairs, and projecting it on the closest histone surface atom, along the direction locally perpendicular to the superhelical axis.

Figure 6a shows the variation of *ζ* as a function of simulation time, at constant pulling force. It can be seen that the DNA broken end detaches from the histone surface in large steps (red segments), during which the internal energy builds up until some kind of barrier is overcome; the final stage, indicated by the blue segment, is the complete detachment of the pulled DSB end after *t*=1.1 ns, in which the free end is simply drifting at the constant speed of about 2 m/s (later on dropping to 1 m/s).

**Figure 6.**
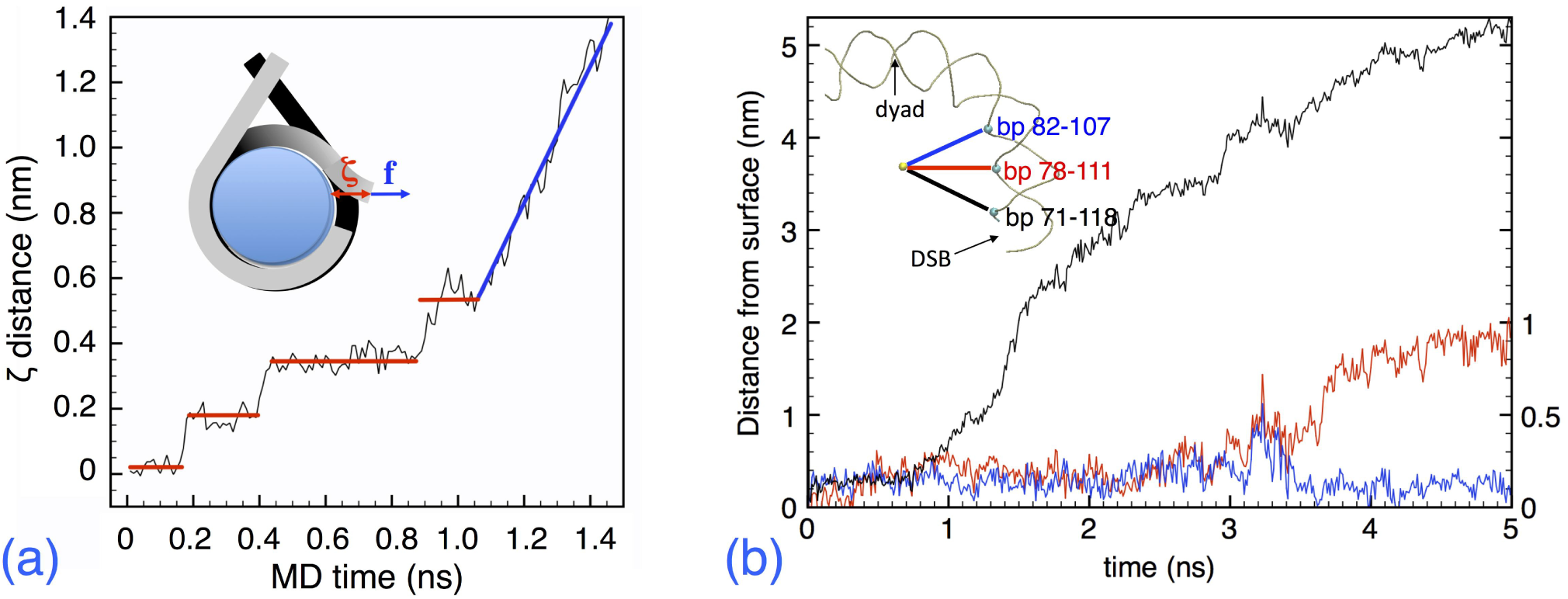
**(a)** Plot of the reaction coordinate *ζ* (distance of the DNA axis center at the DSB site from the closest histone surface) as a function of the MD simulation time, for a pulling simulation at constant force. The first 1.5 ns are shown. The inset schematic shows the definition of the *ζ* distance of the broken DNA end (red arrow), and the direction of the applied force vector (blue arrow), parallel to the *x* axis and perpendicular to the superhelical axis of DNA. **(b)** Plot of the detachment of DNA portions from the histone core surface, from 0 to 5 ns of MD simulation at constant force and 310K. The three traces (black=left ordinates; blue/red=right ordinates) represent the distance to the surface of the three P atoms indicated in the scheme on the upper-left corner, in which a quarter of turn of the DNA comprised between the dyad and the DSB is sketched. The yellow sphere is the geometric center of the nucleosome.

During the final stage of the pulling simulation, the DNA is forcefully unwrapped from the histone core, as it can be seen in Fig 6b. Here we show the distance from the core surface of three P atoms facing the histones, belonging to the bp 71-118 (contact site close to the DSB), 78-111 (middle site) and 82-107 (next contact site). The first contact site is detached in the interval t=1.-1.5 ns, as indicated by the black trace that follows the distance from the surface of of the P71 backbone phosphor. Then, under the continued pulling of the DSB end, also the P111 comes off, at t >3 ns (red trace); however, it may be noticed that this event is ”cooperative”, the P82 (blue trace) following the instantaneous opening of P111 at *t*=3.-3.4 ns, and then falling back into position, after which P111 is definitely ”peeled off” the histone surface.

From this force-pulling simulation we can calculate the free energy profile of the barriers, which characterize the binding of the DNA end to the histone core surface. The potential of mean force (PMF,^39^) is a method to extract the free energy difference ΔA from a sequence of configurations, biased along a reaction coordinate *λ* that brings the system from a state *a* to a state *b*, by estimating the force *f*_*λ*_ necessary to hold the system at each different value of *λ*:

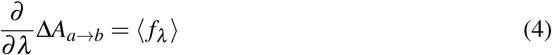

where 〈…〉_*λ*_ means averaging over *a* < *λ* < *b*. In our case, the reaction coordinate *λ* is just the distance *ζ* defined above; the states *a*, *b* respectively represent the initial configuration at *ζ*=0, with the DSB end still attached to the histone surface, and the final configuration with the end detached, at *ζ* ∼5 nm and *t* =1.1 ns. Since the reaction coordinate is arbitrarily chosen, additional care must be taken to allow the system to “explore” as much as possible the nearby configurations, to increase the statistical sampling of possible intermediate states between *a* and *b.* This can be achieved by the so-called The “umbrella sampling” technique,^40^ is used to obtain the PMF at discrete values of *ζ*, and consisting in biasing the true molecular potential with an additional harmonic potential *U*′(*ζ*) at each point along the reaction coordinate, which allows the system to sample configurations in a small parabolic well around *ζ*. The probability of finding the system at *ζ* is now biased, *P*′(*ζ*), and the unbiased estimate of *A* is *A*(*ζ*) = −*k*_*B*_*T*ln*P*′(*ζ*) − *U*′(*ζ*) + *c*, with *c* an undetermined constant that disappears when computing free-energy differences Δ*A*. Finally, the discrete values of *A*(*ζ*) between *a* and *b* are connected by the weighted-histogram method.^41, 42^ We extracted 100 configurations from the force-pulling simulation, spaced by 50 ps in the first 5 ns of the trajectory (corresponding to about 0.5 Å spacing along the reaction coordinate *ζ*=0 to *ζ*∼5 nm); each configuration was equilibrated for 2 ns at 310 K under constant-{*NVT*}, while biased with a harmonic “umbrella” potential of variable strength, progressively reduced to zero to obtain the unbiased limit. The force probability distribution of the fluctuating DSB free-end at each value of *ζ* was reconstructed with the weighted-histogram analysis, and the free energy profile thereby extracted is shown in Figure 7.

**Figure 7.**
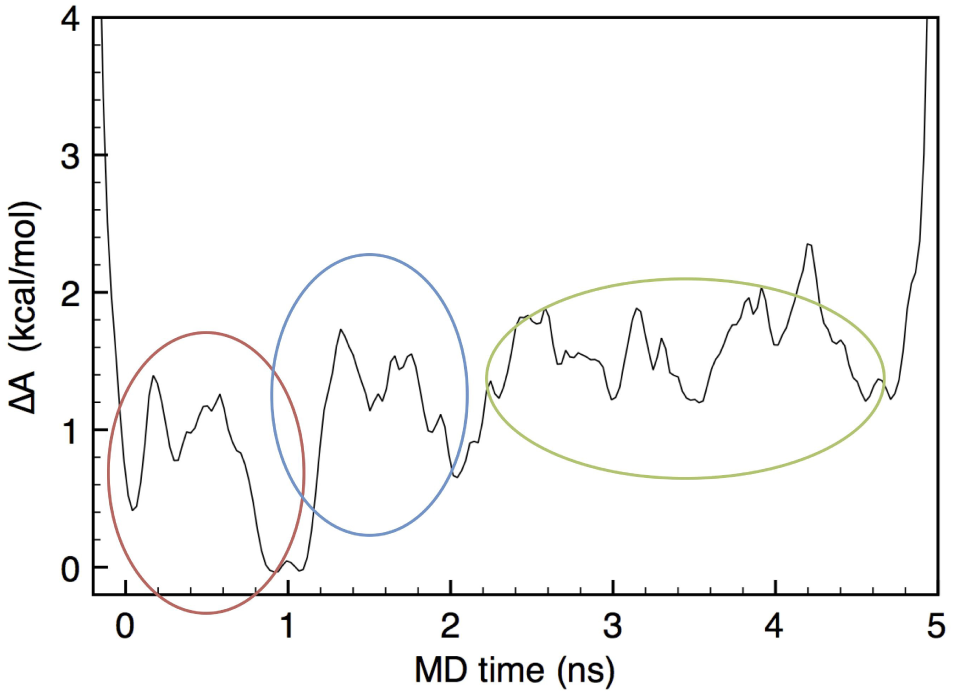
Zero-force extrapolated free-energy profile as a function of MD time, during the force-pulling simulation at T=310 K. The red, blue and green regions refer to the sequence of energy barriers for the DNA-histone detachment events, as described in the text.

From the noisy profile, a few features can be identified. The red circle defines the first barriers to the detachment of the DSB ends, corresponding to the red steps in Fig 6a; such barriers are quite small (<1 kcal/mol), and strongly dependent on the choice of the point of application of the pulling force. The blue circle identifies the free-energy barrier for the detachment of the first contact at P71, about Δ*A*=1.8 kcal/mol or 3 *k*_*B*_*T*; this does not represent a very large value, and should correspond to a ∼5% Boltzmann probability of spontaneous detachment at T=310 K. The green circle roughly identifies the cooperative events leading to the detachment of P111, between 2 and 4 ns, with a sequence of Δ*A* overall not larger than 2-3 *k*_*B*_*T*. Further detachment events were not observed, with the above values of pulling force; in particular, P82 remains in place even at larger deformations of the DSB free end, because of the H3 histone tail acting as a sort of brace that maintains the DNA firmly in place about that position. Much larger forces, or cooperative events of histone tail fluctuation, likely involving other nuclear proteins, seem to be necessary to pull the free DSB end further beyond the limits observed in the present simulations.

### Internal stress relaxation and DSB structure

In the last part of our study, we turn our attention to the internal relaxation dynamics of the nucleosome including a broken DNA. To demonstrate what it is meant by “internal relaxation”, we take two configurations along the final trajectory of the force pulling simulation of the M1 model described in Fig 6, C180 and C250, respectively extracted at times *t*=1.8 and 2.5 ns, well beyond the detachment stage that ends at 1.1 ns in the figure. Each of these two configurations is used as initial structure for a MD simulation, and is then equilibrated and relaxed at 310 K and constant-{*NVT*}, without any external forces applied. The results of these two MD simulations are displayed in Suppl. Fig. 5: starting from the two different initial conditions, after 40 ns the C180 tends to fold back into the initial M1 configuration, while C250 straightens out and increases its distance from the histone core. Notably, the C180 remains in a slightly open state, because of the free energy barrier to detachment that now has to be overcome in reverse, the low temperature not helping in accelerating this last barrier crossing. However, the important observation in both cases is that the folding back, or the straightening out, are driven entirely by the competition between the residual attraction between DNA and proteins (a “chemical” force), and the relaxation of internal constraints (mainly bending and torsion, therefore an “elastic” force). The role of internal forces can be clearly understood by looking at the distribution of mechanical stress, which is a measure of the elastic energy accumulated by the bending and torsion of DNA while wrapping around the histones, and that is ready to be released if the structural constraints are softened, as it could be the case of a DSB cutting the DNA sequence.

Mapping the classical mechanics of a system of point particles to a continuous stress field can be understood in terms of the statistical mechanics framework of atomic force decomposition, based on Liouville’s theorem.^43^ In a particle-based model, the atomic-level stress at an atom *i* is usually defined as the sum of a kinetic and a potential part:^44^

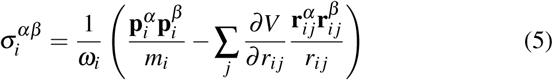

where *m*_*i*_, *ω*_*i*_ and **p**_*i*_ are the mass, volume and kinetic momentum of atom *i*, **r**_*ij*_ is the vector distance between atom *i* and any atom *j* within its interaction range, *r*_*ij*_ = |**r**_*ij*_| = |**r**_*i*_ − **r**_*j*_| its modulus, and *V* = *V*(**r**_1_, …**r**_*N*_) is the interatomic potential comprising all the different force terms; Greek indices *α*, *β* run cyclically over the Cartesian components *x, y, z* of vectors, thereby defining σ_*αβ*_ as a 3×3 tensor. The shortcomings of this definition have been repeatedly underscored.^44–47^ notably in the presence of discontinuities, disorder and defects at the atomic scale, as well as its lack of conservation of linear momentum.^48^

On the other hand, the stress in mechanics is very properly defined as a continuous field, while the atomic decomposition of Eq 5 is based on a not well defined ”atomic volume”. A continuous definition of the (potential part of) stress from the tensor product of force⊗distance at each point in space, σ(**x**), rather than at the atomic sites, is the following:^43^

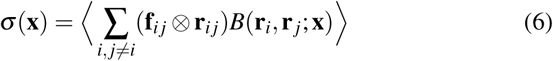

the force vector **f**_*ij*_ = (*∂V* / *∂r*_*ij*_)**r**_*ij*_/**r**_*ij*_ being represented as a sum over the pairwise *i* − *j* interactions, and 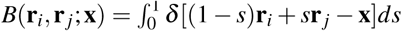 is a continuous bond function,^49^ describing the smearing of the interatomic forces over the space **x** surrounding the interaction region **r**_*i*_…**r**_*j*_.

However, it is well recognized that such a mapping is not unique, particularly in the presence of complex force-fields.^44, 50^ The major ambiguity comes from the non-unique decomposition of the interatomic forces from many-body potentials into pairwise terms, leading to definitions of molecular stress field that can violate conservation of linear momentum, and eventually of angular momentum if molecular chirality is also present. Recent developments in this area led to alternative geometric derivations of the microscopic stress,^33, 51^ based on the invariance of the free energy with respect to surface deformations,^52, 53^ instead of the classical invariance of momentum. The so-called CCFD scheme,^32, 33^ incorporated in a special-purpose version of the GROMACS code, ensures conservation of both linear and angular momentum under a generic stress-induced transformation (diffeomorphism). The continuous stress field components are defined in terms of the variation of the system free energy *δA* with respect to a variation of the metric *δ***g**(**x**):

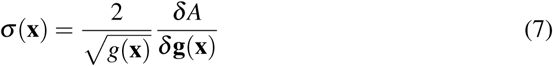

with:

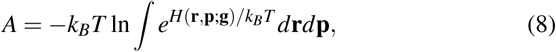

*H*(**r, p; g**) the molecular Hamiltonian describing the kinetic and potential energy, and *g*(**x**) the Jacobian determinant of the metric tensor. The physical meaning of this, quite obscure, differential geometry definition, is (in simplistic terms) that the stress at a material point can be obtained by calculating the change in free energy upon small, arbitrary distortions of the geometry around that point. For a simple pair potential, the expression (7) gives back the Irving-Kirkwood definition (6) plus the kinetic term; however, the CCFD is much more general and provides a proper stress tensor field for an arbitrary intermolecular potential.

The mechanical stress is a meaningful way of representing the distribution of internal forces with respect to a given local direction vector. As a first example of the usefulness of such an information, we represent in Figure 8a the map of the local hydrostatic pressure, that is the trace of the stress tensor *P* = (σ_*xx*_+σ_*yy*_ + σ_*zz*_) /3, averaged over 10 ns of dynamics at T=310 K for the Model O. The stress field in this case is calculated by subtracting from the total stress, the contributions of the isolated DNA plus the isolated histone core, so as to highlight the differential effect induced by their mutual interaction. All continuous stress values here and in the foregoing are spatially-averaged in voxels of (0.1)^3^ nm^3^; we checked that an eightfold denser mesh, of (0.05)^3^ nm^3^, does not add relevant details to the (rather noisy) stress profiles. In the Figure, two isosurfaces corresponding to the values ±70 MPa are shown in red and blue, respectively. Apart from random fluctuations, it can be clearly appreciated that the histone core surface in contact with the DNA is under compressive stress (negative *P),* while the inner DNA surface in contact with the core is under tensile stress (positive *P):* this would be the effect of a stretched elastic band, gently squeezing a compressible body.

**Figure 8.**
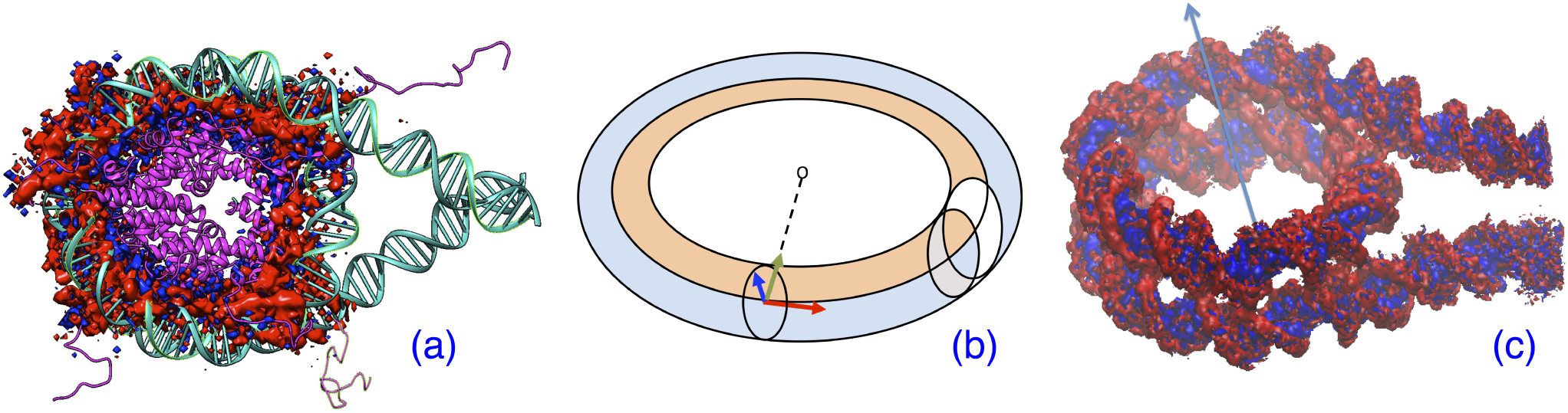
**(a)** Map of the local hydrostatic pressure (trace of the stress tensor) for the nucleosome model O, averaged over 10 ns of MD at 310K. Two isosurfaces at the reference values of +70 MPa (red) and −70 MPa (blue) are shown. The DNA and histone proteins are represented as green and purple ribbons, respectively, with the superhelical axis normal to the figure. **(b)** A schematic cross section of the circular bent tube describing the DNA geometry in the nucleosome. The centerline is the neutral axis. At any point along the neutral axis, a local reference frame can be defined, by introducing the three unit vectors: normal **n** (blue), tangent *τ* (red), and the binomial **b** (green) directed to the center of curvature. The white slice on the right represents one of the tube averaging zones, for computing the tension and twist components of the surface traction. **(c)** Plot of the normal traction s(x) at the DNA surface for Model O at T=310K; stress values averaged over 100 frames spaced by 10 ps. The red and blue isosurfaces represent respectively the values +50 and −50 MPa. To facilitate the view, the model is slightly tilted with respect to the central superhelical axis (blue arrow), and depth cueing is added.

An intuitive way of looking at the stress as a ”projected force” is through the (outer or inner) surface traction vector, **T**(**x**) = σ(**x**) · **n**, with **n** the unit vector normal to the surface at point **x** (see the local reference frame {**n**, *τ*, **b**} in Fig 8b), and the component of the surface traction normal to the surface (a scalar):

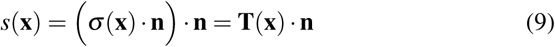

Positive and negative values of *s*(**x**) indicate, respectively, an internal force locally directed outwards or inwards with respect to the surface at the point **x.** In Figure 8c the normal traction is represented on the surface of the DNA wrapped around the histone core in Model O. We show two representative values of positive and negative surface traction, at ±50 MPa, Despite the sizeable noise, the overall distribution of surface traction is clearly evident: the ridges of the phosphate backbone are under outward traction, while the grooves experience an inward directed internal force. However, such internal tensions normal to the DNA surface are perfectly balanced and do not create any net mechanical force, their integral summing to zero. Also, the off-diagonal components of the stress are on average perfectly symmetric, ∑_*α*<*β*_(σ_*αβ*_ − σ_*βα*_) < 10^−4^. Whether such an alternating state of tension and compression perpendicular to the surface could be correlated to some biologically relevant functions is now a matter for investigation.

The traction vector **T**(**x**) contains a great deal of information on the state of internal tension, compression, and torsion, of a complex structure like the DNA in the nucleosome.^2^ The portion of DNA wrapped around the histone core is forced to bend into nearly two full circles of diameter about 8 nm, a size much shorter than the persistence length of free DNA, *ξ*_*p*_ ≃50 nm. Therefore, the DNA “tube” is here constrained in a geometry from which it should rather escape into a more straight structure, whenever possible, under the relaxation of internal forces. The state of tension and compression of a bent tube is described by a different projection of the traction vector, in this case along the unit vector *τ* locally tangent to the continuous line sweeping the center of the tube:

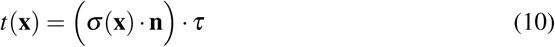

(Note that this is a shearing component of the stress; a pressure component can also be calculated, by replacing σ · **n** with σ · *τ* in the (…). Both projections display the same general behavior, as a function of **x,** however the latter is typically more noisy in the present case.)

Notably, a bent tube will experience a stretching (tensile) force in the half tube which lies outside the centerline with respect to the center of curvature, and a compressive force in the half lying inside the centerline, as shown in blue/orange in Figure 8b. The internal force is zero along the centerline itself, because of this called the ”neutral axis” (also the helical axis of DNA). We computed the line tension *t*(**x**) all along the curved DNA pathlength, by averaging over slices of width 0.5 nm (see for example the white slice in Fig 8b), and by integrating separately over the inner and outer regions (orange and blue in the Figure). Each slice is centered at the midpoint between the two P atoms of each base-pair, therefore adjacent slices have some overlap, to provide a smoother profile of the (very noisy) signal. Figure 9a shows the profile of the line tension along the DNA in nucleosome model O, for the bp 21 to 167 that wrap around the histone core; the upper line corresponds to the inner (orange) part of the DNA ”tube”, and is in compression; the lower line corresponds to the outer (blue) part, and is under tension. It can be seen that both the tension and compression sides display 14 broader maxima, likely corresponding to the contact regions of DNA with the histone core surface. Notably, the maxima of the tension side roughly correspond to the minima of the compression side, and viceversa, the maximum compression and minimum tension being nearly aligned with each contact site. Note that for a tube bent into a perfect circle (torus) with constant radius of curvature, one should observe constant values of tension and compression all around the circle, therefore the alternating minima and maxima in the stress indirectly demonstrate the extra curvature at the DNA-histone contact sites.

**Figure 9.**
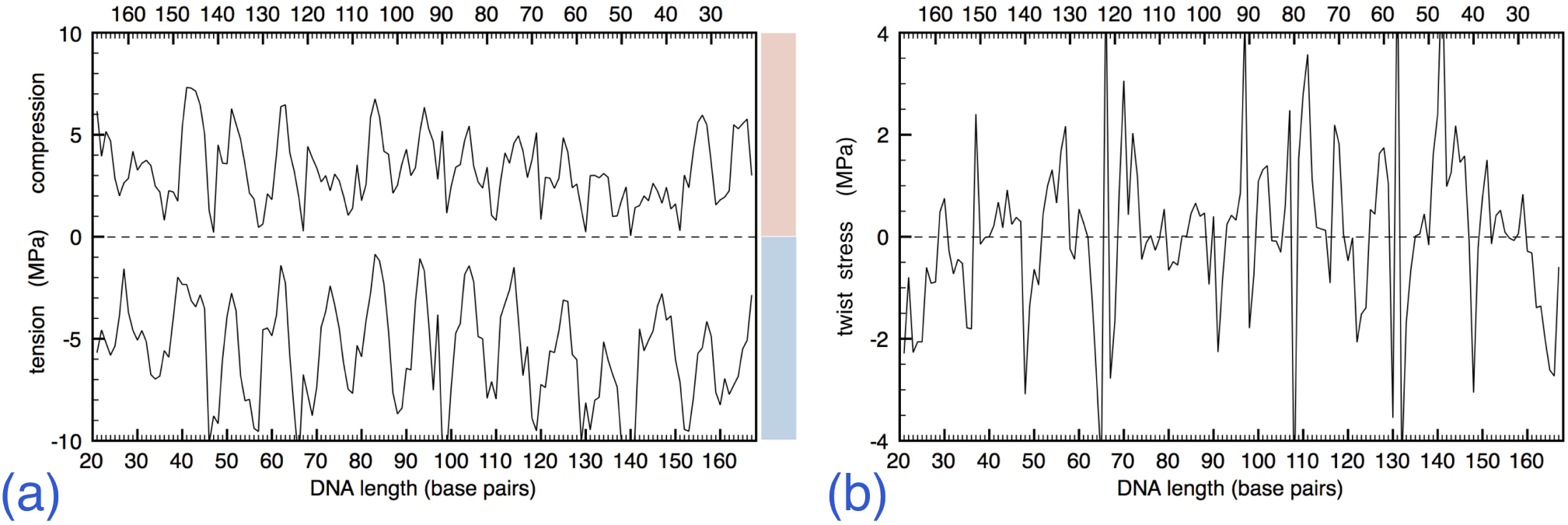
**(a)** Plot of the tension/compression force *t*(**x**) along the DNA model O. Numbers along the upper and lower ordinates indicate the DNA bases in the two strands; only the bp 21-167 wrapped around the histone core are shown; color bars on the right refer to the “tube” regions in Fig 8b. (b) Plot of the twist force *w*(**x**) along the DNA model O.

The twist stress is that part of the internal forces involved in the torsion about the central (neutral) axis of the tube. The DNA double helix is naturally twisted already in its normal B configuration; however, when it is bent in the nucleosome, the twist is necessarily modified with respect to the normal configuration. The twist component is obtained as well from the traction vector, as:

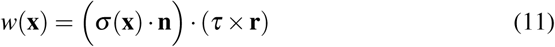

where **r** is the vector from the neutral axis to the point **x**, parallel to the local normal to the outer tube surface **n** (see again Fig 8b). Fig 9b displays the twist stress along DNA bp 21-167 of nucleosome model O. The absolute value of stress, fluctuating between about ±4 MPa, is quite smaller compared to the values of line tension or compression; this quantity fluctuates between positive and negative, indicating a force tending to over‐ or under-twist the DNA about its helical axis. One can observe again an alternation between positive and negative values, grouped in 13-15 broad periods; however, alignment with DNA-histone contacts is quite less evident, and sudden, random oscillations between negative and positive values are seen.

Once a DSB breaks the DNA backbone around the nucleosome, such internal forces are going to be relaxed, and compete with the chemical (Van der Waals, electrostatic) forces from the interaction with the histone proteins. Going back to Suppl. Fig 5 for model M1, such a competition is very evident upon comparing the bottom configurations: in C185 the chemical forces overwhelm the internal stress, whereas in C250 the opposite holds, and the DNA ends up straightened out from the DSB site. In Figures 10a and 10b we show for both configurations, the tension profile *t***(x)** along the helical axis of the DNA fragment right after the DSB, bp 72 to 167, i.e., the DSB end that was pulled out under force and subsequently relaxed; we neglect the first few bp immediately next to the DSB, too disordered for such a calculation. Two sets of data are shown in each panel, at the beginning of the relaxation (black lines), and after 40 ns (red lines); stress values are averaged over 100 frames with 10 ps spacing, in either case. In general, the terminal part of the DNA next to the DSB (indicated by a grey-shaded area in the panels) tends to lower values of both line tension and compression, for both configurations, compared to the rest of DNA beyond the dyad (bp 94-94). The data are rather noisy, such a noise being largely numerical (not arising from the thermal average), coming from the difficulty of properly identifying the local reference frame {**n**, *τ*, **b**} at each point **x** along the fluctuating DNA helical axis (centerline). However, it may still be appreciated that for the C185 configuration, Fig 10a, the red lines are at the same values, if not slightly larger, than the black ones: this is a signature of the chemical residual attraction winning over the internal stress, thus tending to fold back the DSB open end into place. On the other hand, the C250, Fig 10b, has red lines approaching a state of zero tension/compression, indicating the release of internal stress, which is responsible for straightening out the DSB end, into a mechanically less-constrained structure.

**Figure 10.**
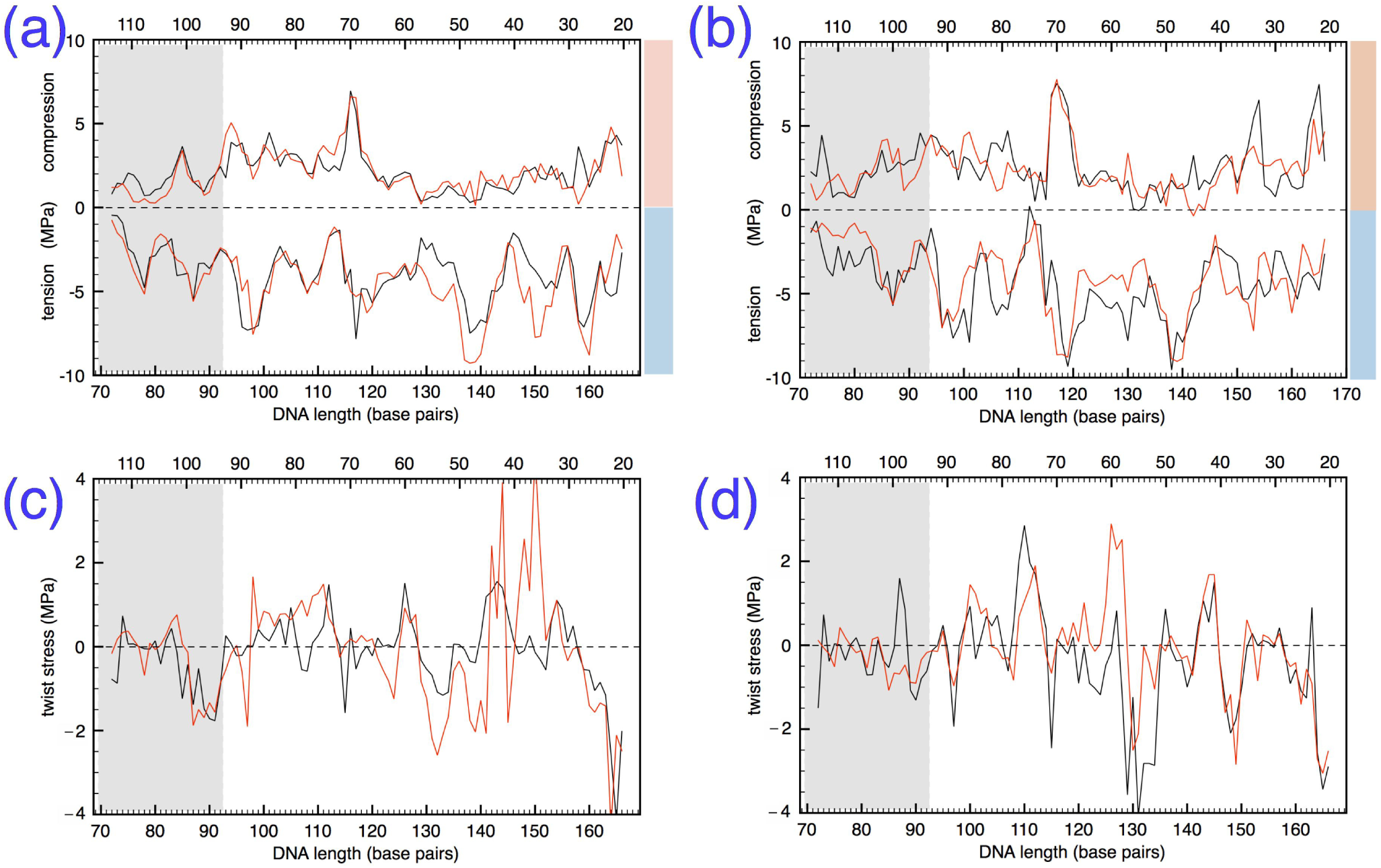
**(a)** Plot of the tension/compression force *t*(**x**) along part of the DNA model M1 configuration C185, at the beginning of the relaxation (black line) and after 40 ns (red line). Numbers along the upper and lower ordinates indicate the DNA bases in the two strands; the DSB is at the extreme left, bp 68-120; the grey rectangle is the main stress relaxation region, comprised between the DSB and the dyad (bp 94-94); color bars on the right refer to the regions in Fig 8b. **(b)** Same as (a), for the M1 configuration C250. **(c)** Plot of the twist force *w***(x)** along the DNA model M1 configuration C185. **(d)** Same as (c), for the Ml configuration C250.

The extra twist stress (positive or negative) also contributes to the internal forces that are going to be relaxed, when the DSB cuts open the DNA, albeit to a much lesser extent, given its absolute value. In Figs.10c and 10d the profile of *w*(**x**) is displayed, under the same conditions of the two panels above for the line-tension/compression. It can be noticed that, also for the twist stress, generally smaller values (≤ 1 MPa in modulus) are seen in the DSB tail comprised in the grey-shaded part. However, the numerical noise does not allow in this case to draw a more firm conclusion, concerning the (likely minor) role of twist stress in the chemical vs. mechanical force competition in the two configurations.

## Discussion

In the present work, we studied by very-large-scale molecular dynamics (MD) simulations the evolution under external force and temperature of double-strand breaks (DSB) in nucleosomal DNA. We collected and analyzed a large amount of raw data (more than 1.5 TBytes, and 5 million CPU hours on two large supercomputers), by running microsecond-long trajectories for 5 different, all-atom models of the experimental 1kx5 nucleosome structure.^22^ The basic model is made up of the canonical 8 histones, plus a 187-bp DNA comprising the 147 bp wrapped around the histone core and 20-bp terminations on each end, and embedded in large boxes of about 80-100,000 water molecules with Na^+^ and Cl^−^ ions at 0.15 M physiological concentration. The pristine nucleosome configuration (model O) was modified, by inserting a DSB at four different positions in the DNA (models M1-M4), and the stability of the resulting structures was compared with model O nucleosome.

A general observation from the microsecond-long trajectories, is that damaged DNA remains well attached to the nucleosome body, without qualitative differences compared to the intact DNA. Only the model M1, in which the DSB is tightly sandwiched between the histone H3 at the tail of histone H2B, displayed a dynamics substantially different from the corresponding region in model O, due to the increased interaction of the broken DSB ends with close-by histone residues; however, also this DSB configuration was stable over the entire observation time scale, which in this case was extended to 1.8 μs. In order to identify the free-energy barriers which maintain the broken DNA attached to the histone core, we carried out steered MD with a pulling force to ”peel off” the free DSB end from the nucleosome; relatively small barriers of the order of 3 *k*_*B*_*T* were identified by umbrella sampling of the reaction coordinate defining the distance of the DSB free end from the histone surface, which seem to point to the possibility of spontaneous detachment at physiological temperatures, likely over longer time scales of hundreds of microseconds to milliseconds. Spontaneous unwrapping of DNA from the nucleosome core has been studied experimentally,^54–56^ because of its relevance in gene regulation and DNA transcription; notably, such experiments were carried out on isolated nucleosomes, with a length of DNA just matching, or barely longer than needed to wrap the histone core (147 to ∼180 bp). In such conditions, spontaneous detachment of the ends was indeed observed over the timescale of hundreds of milliseconds; simulations by coarse-grained MD methods roughly confirm such trends,^57–59^ despite being strongly depending on the empirical parametrization of each different force model. To such experiments it may be objected that the nucleosome constrained in the chromatin could have a rather different mechanics: our molecular stress calculations demonstrate that the circularly bent DNA has a strong internal driving force, from the relaxation of line tension and, to a lesser extent, of twist (torsional) stress. The reason may be found in the persistence length of the free DNA, which is much longer (∼50 nm) than the average radius of curvature in the nucleosome (∼8 nm), and pushes the DNA to regain the straight average conformation on that length scale; the fact that spontaneous fluctuations were observed^55^ both in presence and without binding proteins seems to support this view. In fact, our μ*s*-long MD simulations were carried out with a soft restraining of the DNA linker (20 bp on each end), to simulate the effect of the background chromatin structure, and no fluctuations larger than thermal vibrations were detected for the terminal phosphors; on the other hand, the DSB free end, once extended beyond a distance of about 2.5 nm away from the histone core, tended to regain a straight conformation and detach completely, confirming the importance of stress relaxation as a main driving force.

In conclusion, the results and potential implications of this study can be summarized by the following findings:

1. The closely-cut DSB remain relatively stable over long time scales, and display no sign of disassembly; interaction of DSB ends with histone surfaces and tails is a main factor in damaged-DNA dynamics. DSB configurations close to histone tails in fact displayed a more active internal dynamics, with a participation also from histone fragment fluctuations.
2. The free-energy barriers for detachment of DNA from histones are relatively low, of the order of a few *k*_*B*_*T*, implying that short sections of DNA could spontaneously unwrap over a time scale of > 100 microseconds, from DSB broken ends, or from the linker sections at the nucleosome ends, as observed in some experiments.^54–56^ At the same time, histone tails represent a major steric obstacle for unwrapping of larger DNA sections, notwithstanding the driving force from stress relaxation (3. below).
3. We performed, for the first time, fully consistent molecular stress calculations on the DNA wrapped in a nucleosome; this study revealed the existence of a strong internal driving force for straightening the circularly bent segments, from the relaxation of line-tension and torsional stress. This might be the main force leading to spontaneous unwrapping of DSB cut ends, as well as of nucleosome ends, respectively opening the way to damage-signalling and repair proteins, and to remodelling factors.

## Acknowledgments

Useful discussions about definitions of continuum stress with S. Giordano (IEMN Lille) are acknowledged. We gratefully thank partial funding from the SIRIC OncoLille under project ”ModCel” for 2015 and 2016, and a three-year doctoral grant to F. L. co-funded by the President of Lille-I University and the Government of the Region Nord-Pas de Calais (now Hauts-de-France). Computer resources provided by the CINES and IDRIS French Supercomputing Centres, under grants a0020707225 and x2016/077225, and by generous extensions thereof.

## Author contributions

F.C. and R.B. designed the experimental work. F.C. and F.L. performed the computer simulations. F.C. drafted the paper. All authors contributed to the final written document.

## Supplemental data

Supplemental Data include five figures and can be found with this article online at http://www....

## SUPPLEMENTAL DATA

**Supplementary Figure 1.**
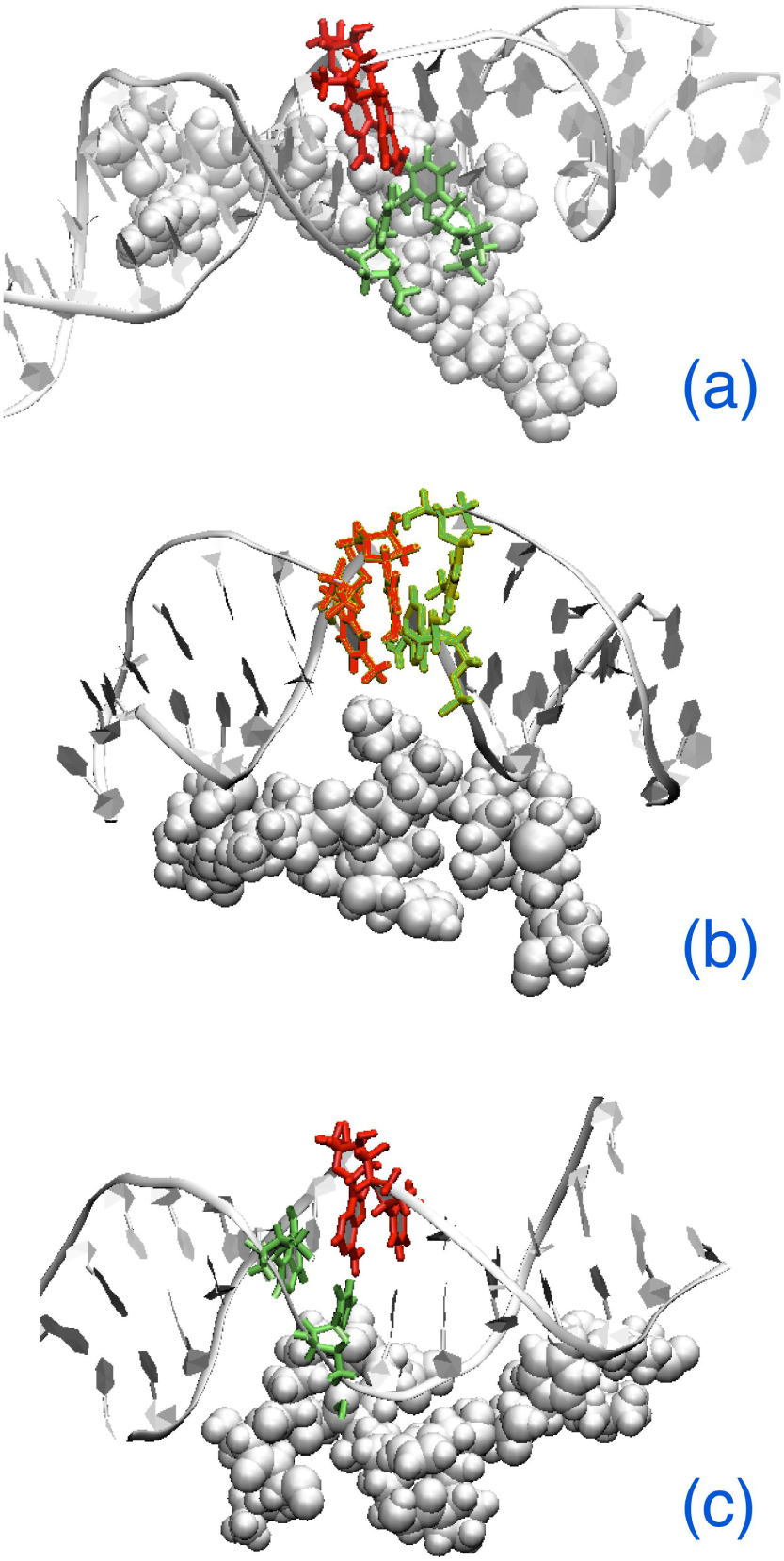
**Snapshots of MD simulations of the M2-M4 models of DSB,** after 1 μs of dynamics atT=350 K. **(a)** Model M2, with DSB at the outer non-contact site, showing the central A74… T114 bp still well bonded. Grey spheres represent a portion of the H3 histone flanking the defect. **(b)** Model M3, with DSB at the dyad. Grey spheres represent a portion of the H3 tail. **(c)** Model M4, with DSB at the entry point of nucleosomal DNA. Grey spheres represent a portion of the H3 tail close to the break, which has folded into a double *α*-helix.

**Supplementary Figure 2.**
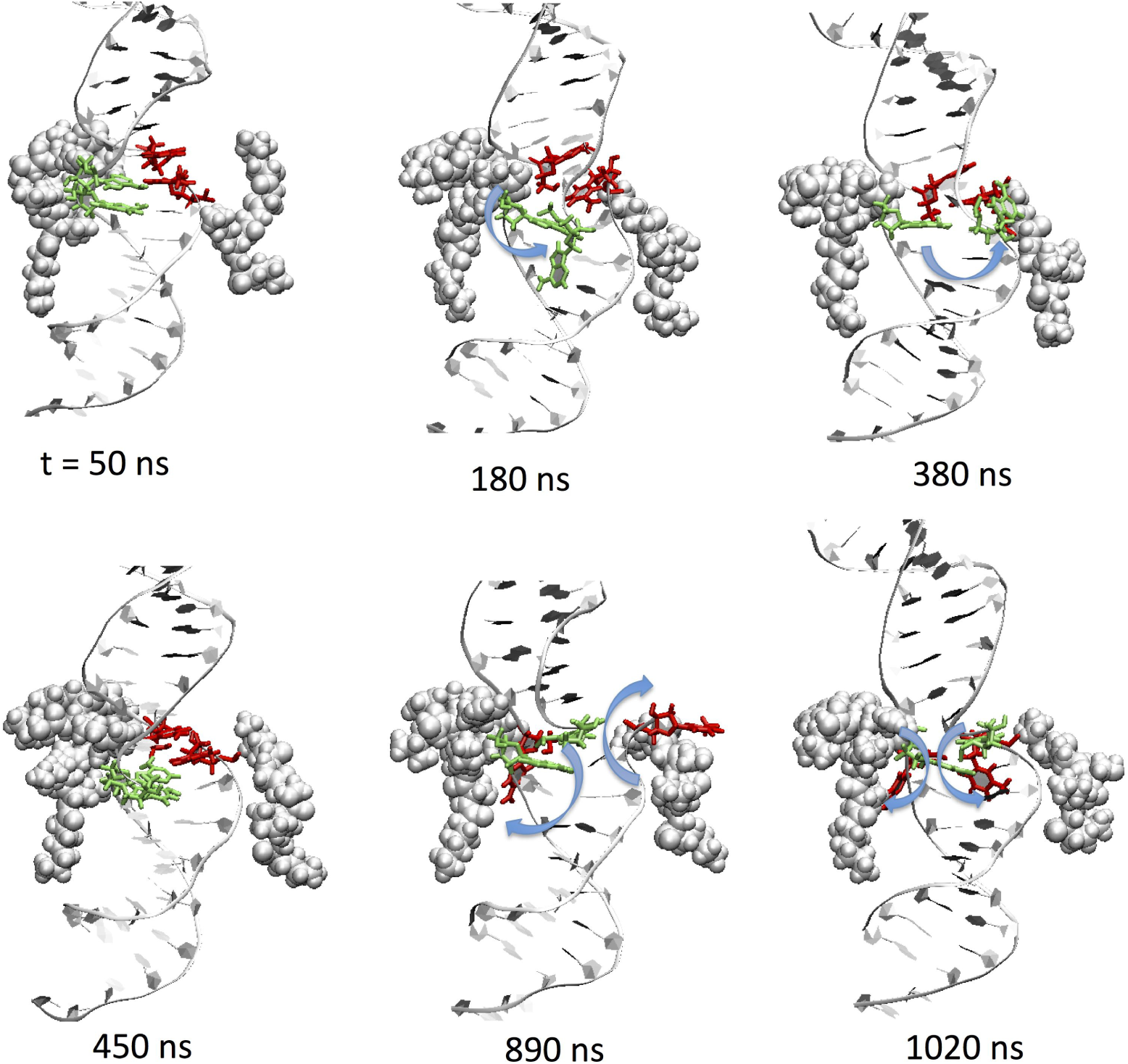
**Time-sequence of the evolution of the DNA-histone contacts** for the DSB at the M1 position. The groups in grey VdW-spheres are the Leu82-Arg83-Phe84-Gln85 of H3, Lys77-Arg78-Lys79-Thr80 of H4 (left side); and Lys9-Gly10-Ser11-Lys12-Lys13, Lys24-Lys25-Arg26 of H2B (right side). DNA bp around the DSB are colored red-green according to the scheme of Fig.1b.

**Supplementary Figure 3.**
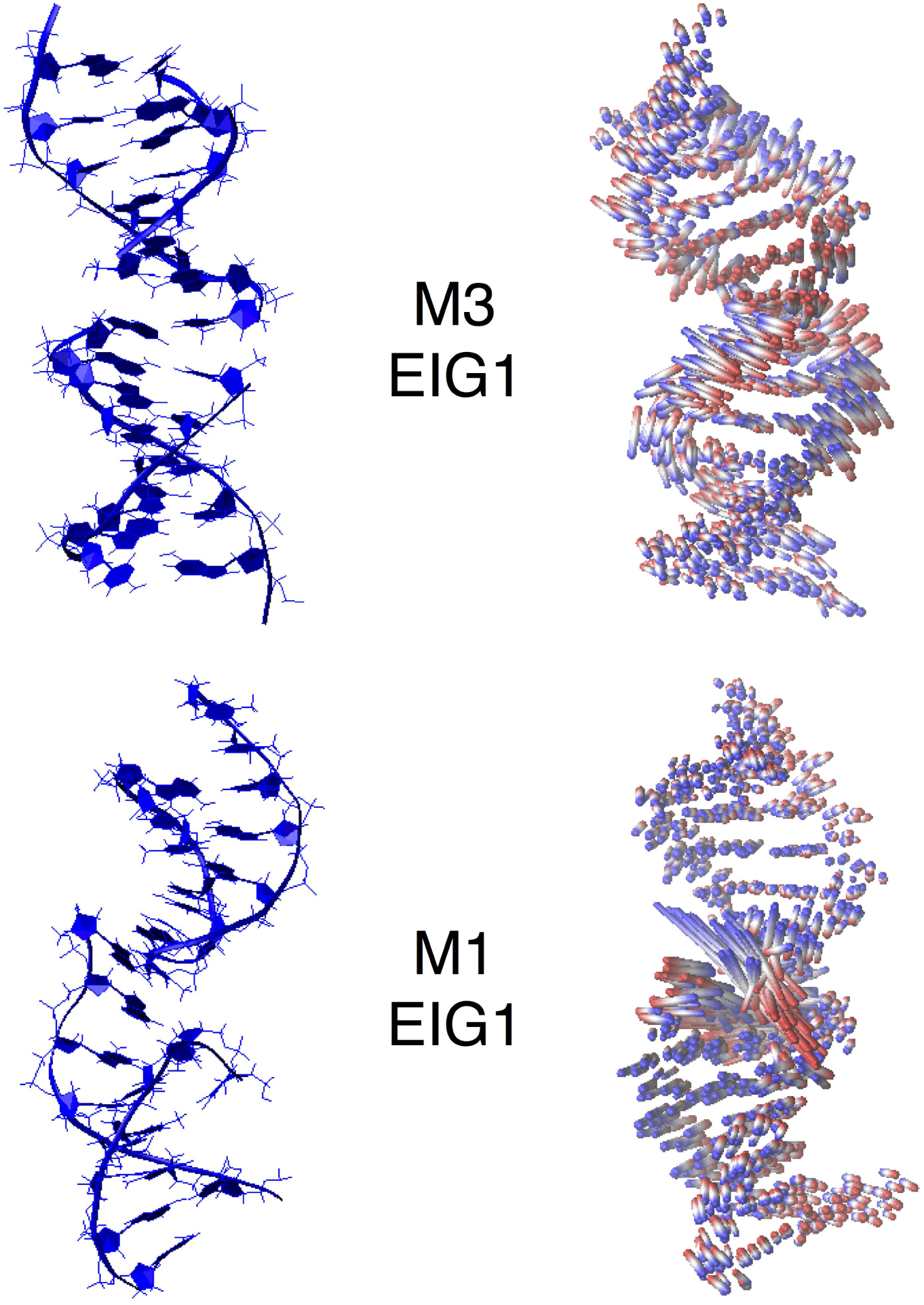
**Configurations associated with first eigenvector of the covariance matrix.** On the left, representative configurations of the DNA fragments close to the DSB, in models M3 (above) and M1 (below). On the right, simultaneous plot of the configurations spanned by the principal motions associated with the first eigenvector, for each model. DNA fragments are aligned with their main axis vertical, the DSB being at the center. The superimposed frames are colored from blue to red, the ordering reflects a virtual motion spanning between the eigenvector extremes. A long stick spanning between the two colors identifies a large motion of the corresponding atom; a shorter stick identifies a local oscillation, of smaller amplitude.

**Supplementary Figure 4.**
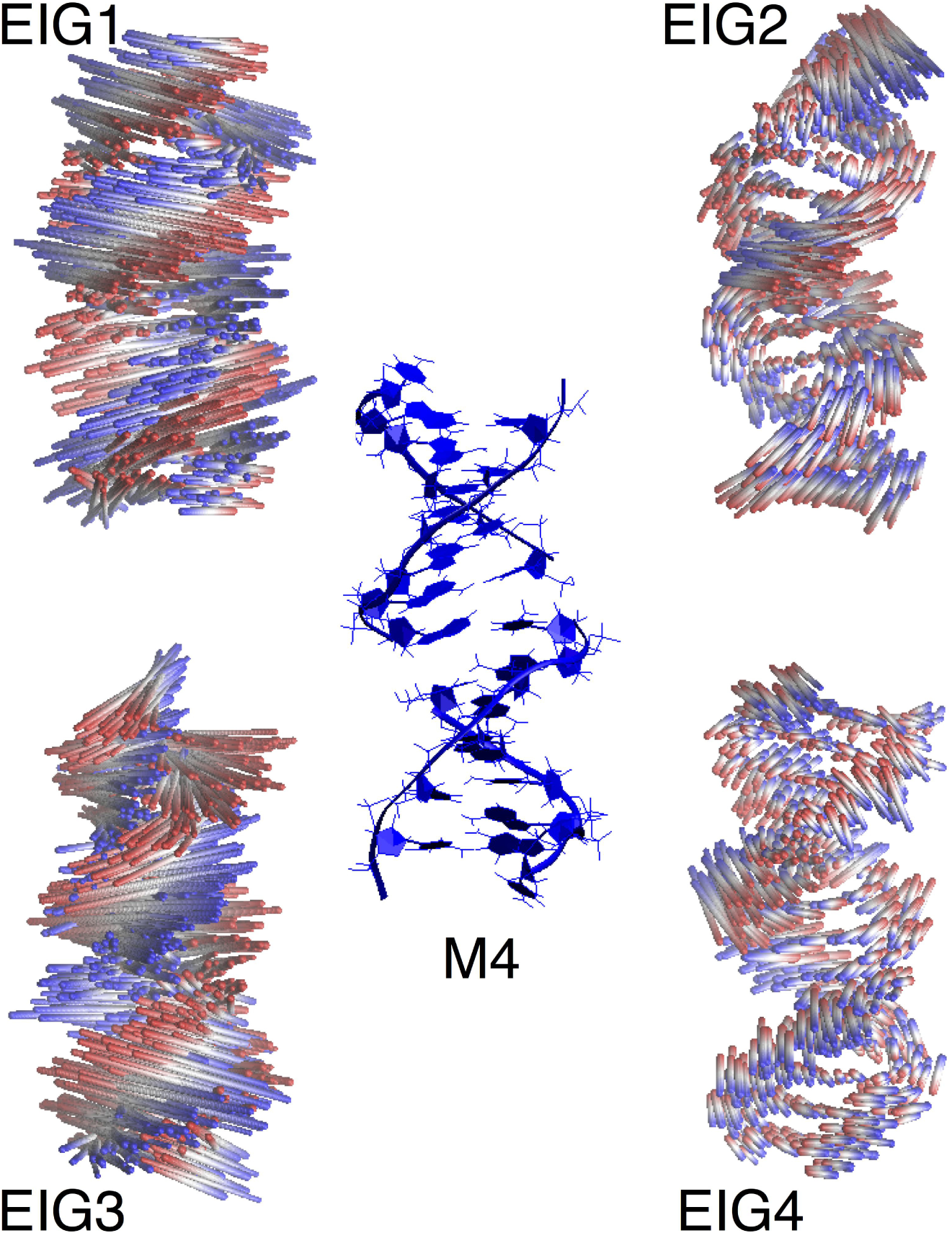
**Configurations associated with eigenvector 1-4 of DSB model 4.** Simultaneous plot of the configurations spanned by the principal motions associated with the eigenvectors 1-4, for the DNA fragment close to the DSB M4 (represented in the central panel). The superimposed frames are colored from blue to red, the ordering reflects a virtual motion spanning between the eigenvector extremes. Also in this case, a long stick spanning between the two colors identifies a large motion of the corresponding atom; a shorter stick identifies a local oscillation, of smaller amplitude. It can be readily appreciated that eigenvectors 1 and 3 correspond to a coordinated, twisting motion of the entire fragment, while eigenvectors 2 and 4 correspond to smaller and less cooperative deformations.

**Supplementary Figure 5.**
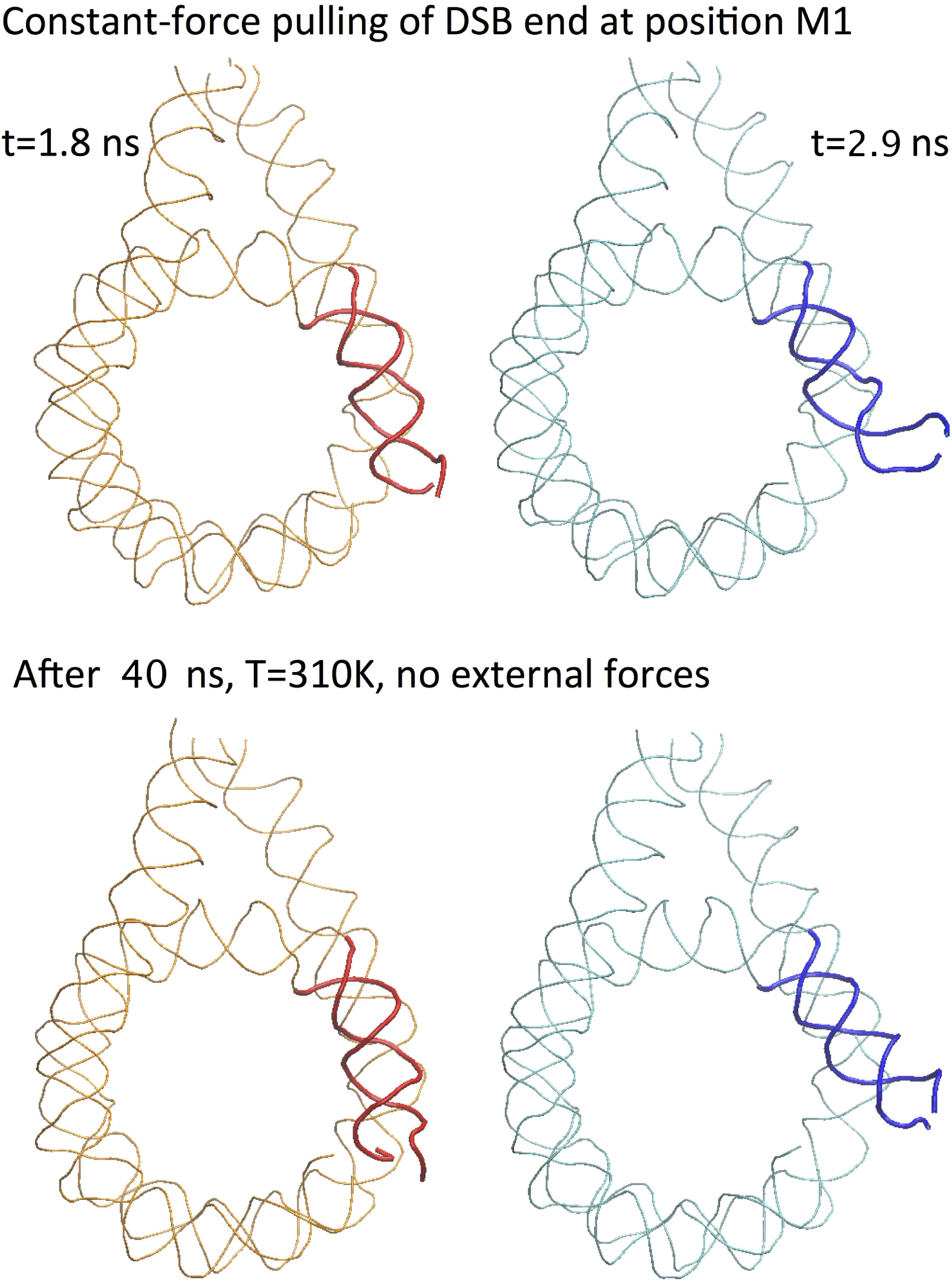
**Stress relaxation in a detached DSB.** Two configurations extracted from the force-pulling MD simulation of DSB at position M1, at *t*=1.8 ns (C180, left, red ribbons) and *t*=2.9 ns (C290, right, blue ribbons). The pull force was applied only at the *C*′-P atoms of the 2 last bp on the upper end of the DSB. The terminal portions of the pulled DSB end are highlighted as a thicker tube, for clarity. Row above: the two configurations at the start of the relaxation. Row below: the two configurations after 40 ns of MD equilibration/relaxation at T=310 K without any external forces applied.

## Footnotes

1 The – symbol indicates the break site along each backbone, the … indicate the central interacting base pair.

2 Of course, a tensor can be meaningfully projected onto any direction vector, the choice of a particular projection being just a matter of convenience. In the present case, the “bent tube” structure of nucleosomal DNA makes it interesting to consider the stress projected onto its “tubular” surface.

